# COP9 complex maintains neuroblast growth and proliferation by regulating Akt/mTor pathway

**DOI:** 10.64898/2026.07.23.740252

**Authors:** Nandan Jayaram, Priyanka Pandey, Syed Arijmand, Deepa Balasubramanian, Manish Jaiswal, Sonal Nagarkar-Jaiswal

## Abstract

The nervous system consists of the brain and associated structures that are replete with highly specialised neurons and glia. Central to the development of the brain are Neural Stem Cells, which self-renew to maintain their number and also give rise to differentiated progeny. To identify genes involved in the maintenance of *Drosophila* neural stem cells, neuroblasts, we performed a protein expression screen followed by a protein knockdown screen using deGradFP system. Through this, we identified *CSN7*, a COP9 signalosome (CSN) subunit, which is enriched in neuroblasts and essential for neural development. CSN is a highly conserved multi-protein complex that regulates proteasome-mediated protein degradation via modulation of Cullin-RING E3 ligases. We found that loss of *CSN7* and *CSN1b* lead to a decrease in neuroblast size and a reduced mitotic index. Our results show that *CSN7/CSN1b* regulates Akt-TOR signalling in the developing larval brain. Furthermore, we found that this regulation is mediated by Cul1. Overall, our work describes a hitherto undescribed role for the components of the CSN complex in neural development.

## Introduction

The nervous system consists of the brain and associated structures that are replete with highly specialized neurons and glia. Central to the development of such a complex and highly organized system are multi-potent stem cells known as neural stem cells (NSCs). During development, NSCs self-renew to maintain a stem cell pool and give rise to daughter cells that differentiate and generate neurons (Morrison *et al*, 1997). The precise regulation of NSC self-renewal and differentiation is pivotal in neurogenesis. *Drosophila* NSCs, the neuroblast (NB), serve as an excellent model system to study neural development and a valuable tool in deciphering the basic mechanisms that underlie NSC maintenance and self-renewal. Based on lineage, the *Drosophila* central nervous system has two types of NBs-type I and type II-both divide asymmetrically and generate neurons and glial cells (Boone & Doe, 2008). The Type I NB produces a daughter NB and a ganglion mother cell (GMC), which divides symmetrically, differentiates, and gives rise to two neurons or glial cells (Karcavich & Doe, 2005, Boone & Doe, 2008). The type II NBs generate one daughter NB and an intermediate neural precursor (INP). INPs undergo several rounds of asymmetric divisions and in each round generate one INP and two neurons/glia. The prerequisite to asymmetric division is the establishment of an axis of polarity along the apical-basal axis (Knoblich, 2008). In the early embryo, when NBs delaminate from the neuroepithelium, the cells inherit the apical-basal axis from the overlying epithelium (Rusan & Peifer, 2007). The apical region of the NB is defined by the presence of a set of proteins, primarily the Par complex (Bazooka, Par- and aPKC) (Boone & Doe, 2008) (Ohshiro *et al*, 2000), these then direct the localization of the cell-fate determinants; Brain tumour (Brat), Prospero (Pros) and Numb to the basal cortex and their adaptor proteins Miranda (Mira) for Pros and, Brat and Partner-of-Numb (Pon) for Numb to the basal cortex (Chia *et al*, 2008; Ikeshima-Kataoka *et al*, 1997). Upon division, the cell-fate determinants segregate to GMC. Within the GMC, the adaptor proteins are degraded, releasing their cargo to promote differentiation. After each round of asymmetric division, NB re-grow to their original size; this re-growth is prerequisite for the next round of division.

The COP9 signalosome (CSN) complex is a highly conserved complex found in eukaryotic cells, playing crucial roles in regulating various cellular processes. It is a multi-protein complex thought to canonically consist of 8 subunits (termed CSN1-8) (Wei & Deng, 2003); however, recent work suggests the presence of a 9th subunit (CSN-AP) (Füzesi-Levi *et al*, 2020). CSN serves as a multifunctional regulator, involved in diverse cellular activities such as protein degradation (Wei & Deng, 2003), cell cycle progression (Björklund *et al*, 2006; Menon *et al*, 2007; Liu *et al*, 2013), DNA repair (Meir *et al*, 2015), and transcriptional regulation (Chamovitz, 2009); it orchestrates these processes by modulating the levels and activities of key proteins. The primary function of the complex is the regulation of a class of E3 ubiquitin ligases known as the Cullin-RING ligases (CRL) that require the covalent addition of Nedd8, a ubiquitin-like protein, for functionality (Cope *et al*, 2002; Hotton & Callis, 2008). CSN possesses iso-peptidase activity that cleaves Nedd8 from E3 ligases (deneddylation), thereby modulating their activity and substrate specificity (Cope *et al*, 2002). The catalytic activity of the CSN complex is contributed by CSN5, which contains the requisite JAMM-MPN motif; however, the whole complex is required for the function (Ambroggio *et al*, 2003; Wei & Deng, 2003). In addition to its role in regulating proteolysis, CSN is also known to recruit kinases that phosphorylate cell-cycle regulators such as p53 and target its degradation (Schwechheimer, 2004). CSN is also known to be involved in the regulation of cell-cycle regulators such as p27 (Yang *et al*, 2002), p53 (Zhang *et al*, 2008; Lykke-Andersen *et al*, 2003), and Rb proteins (Ullah *et al*, 2007). Studies on human T-cells have shown that CSN binds to and possibly regulates Merlin, a known upstream regulator of the Hippo pathway that is reported to be involved in maintenance of quiescence (Stotland *et al*, 2012). Some of the COP9 complex subunits have also been shown to function independently of the holocomplex. For example, CSN1 and CSN2 are required for DNA damage response and progression through S phase in S. pombe (Suh *et al*, 2002). In *Drosophila*, all of the COP9 subunits are conserved (Freilich *et al*, 1999) though only CSN4, CSN5, and CSN8 have been studied in detail, and they are known to be required for larval development and oogenesis (Freilich *et al*, 1999; Oron *et al*, 2002; Djagaeva & Doronkin, 2009; Doronkin *et al*, 2002; Oron *et al*, 2007; Doronkin *et al*, 2003). Precursory studies on CSN7 have shown that it binds to DNA loci throughout the genome and has been demonstrated to be involved in wing disc development and oogenesis (Singer *et al*, 2014, Doronkin *et al*, 2003). Several reports suggest that the CSN subunits may have individual functions independent of the holocomplex. For example, mutations in CSN4 and CSN5 are known to result in patterning defects in *Drosophila*; however, while CSN4 mutants bring about molting defects and melanotic tumors, CSN5 does not, suggesting they act separately in addition to their role in the complex (Oron *et al*, 2002).

In this study, we used MiMIC RMCE lines expressing endogenously EGFP-tagged genes (Nagarkar-Jaiswal *et al*, 2015) (https://flypush.research.bcm.edu/pscreen/rmce/) and performed an expression-based screen followed by a protein knock-down screen using deGradFP (Caussinus & Affolter, 2016) to identify genes that are expressed in NBs. We identified that the highly conserved multi-subunit COP9 Signalosome (CSN) complex plays an important role in brain development. The *CSN7-* and *CSN1b-*deficient NBs are significantly smaller in size and have reduced proliferation. Moreover, they have reduced Akt levels and accumulate the active form of Cul1 (neddylated). Further, we show that overexpression of Akt or reduction of *Cul1* in *CSN7-* and *CSN1b-*deficient NBs rescues the NB growth and proliferation defect. Our data suggest that the CSN complex regulates NB growth and proliferation by modulating Akt levels through deneddylation-mediated Cul1 deactivation.

## Results

### COP9 Signalosome complex is necessary for neural development in *Drosophila*

To identify new players involved in NB maintenance, we performed an extensive expression-based screen on fly lines expressing endogenously EGFP-tagged genes (Nagarkar-Jaiswal *et al*, 2015). We screened third-instar larval brains from 200 MiMIC RMCE lines using anti-GFP and anti-Miranda (NB marker) antibodies and selected 30 genes that have enriched GFP expression in NBs for further investigation (Figure S1-A). Next, we knocked down GFP-tagged proteins in NBs using the deGradFP system, which allows conditional knockdown of GFP tagged proteins (Caussinus & Affolter, 2016). The deGradFP protein was expressed using *insc-GAL4, UAS-mCD8GFP/Wor-GAL4* (NB-specific driver), and deGradFP (Figure S1-B). We found that in 13 out of 30 lines, deGradFP-mediated protein knockdown resulted in a small brain phenotype; these are likely to be essential for brain development (Figure S1-C).

Among the 13 candidate genes, we focused on *CSN7*, a subunit of COP9 Signalosome (CSN) complex. Our expression study shows that the CSN7::GFP::CSN7 protein is enriched in NBs (Figure 1-A). To confirm the effects of loss of *CSN7* function on brain development, we combined mutant alleles of *CSN7* (*y*w*; CSN7^Mi07174^*/*CSN7^Mi13910^* and *y*w*;CSN7^M13910^*/*CSN7^T2A>GAL4^*) in hetero-allelic combinations. We observed that these larvae are developmentally delayed and, compared to controls had a severe reduction in brain lobe volume (Figure 1-B). To check that small brain phenotype and larval lethality are due to *CSN7* loss, we generated transgenic lines expressing CSN7::HA protein (Figure S1-D) and performed rescue experiments. We expressed CSN7::HA in *CSN7^Mi13910^*/*CSN7 ^T2A>GAL4^* animals, where expression of CSN7::HA is driven by *CSN7 ^T2A>GAL4^*, an endogenous GAL4 driver, where GAL4 expression is regulated by CSN7’s native regulatory elements (Diao & White, 2012). We found that expression of CSN7::HA rescues the small brain phenotype and larval lethality, indicating that these phenotypes are due to the loss of *CSN7* function (Figure 1-B, iv-v). Further, we proceeded to ascertain whether the entire CSN complex contributes to brain development. For this, we selected *CSN1b* subunit and performed knockdown in the developing brain using *insc-GAL4, UAS-mCD8GFP* driver, and two previously verified RNAi lines against *CSN1b*, GD11099 (VDRC) and TRiP.JF02612 (BDSC). Our results show that loss of *CSN1b* also leads to a decrease in brain size (Figure 1-C), which further confirms that the entire CSN complex is involved in brain development.

**Figure 1:**
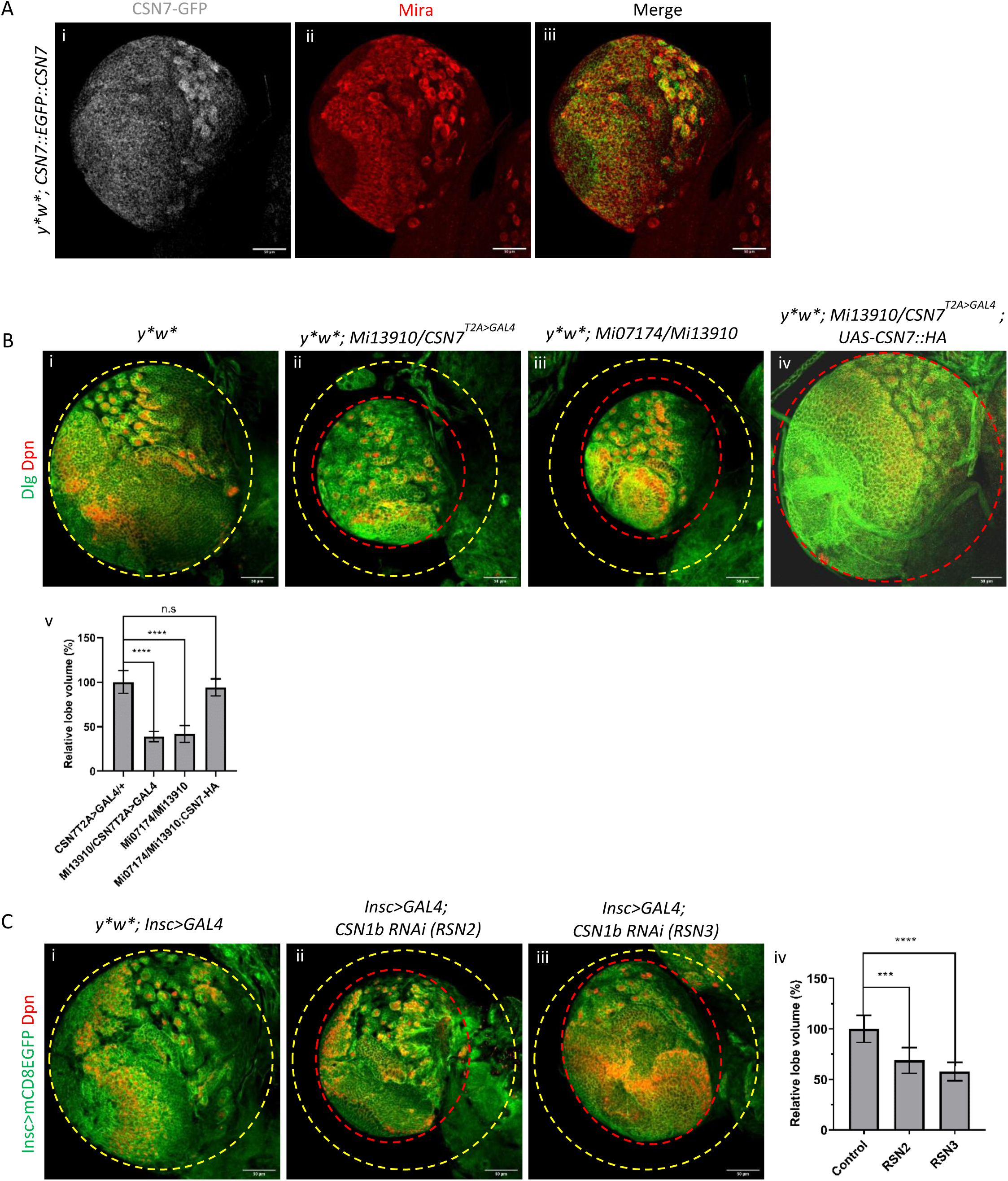
Loss of *CSN7* and *CSN1b* leads to small brain phenotype. (A) Third-instar larval brain lobe expressing *CSN7::EGFP::CSN7* stained with anti-GFP (i), anti-Mira (ii), merged image (iii) showing that *CSN7::EGFP::CSN7* co-localises with Mira in larval neuroblasts. (B) Larval brains (72 hours ALH) from *y*w\** control (i, n=21), *CSN7* mutants *y*w*; Mi13910/CSN7^T2A>GAL4^* (ii, n=24) and *y*w*; Mi07174/Mi13910* (iii, n=24) exhibiting small brain phenotype and *y*w*; Mi13910/CSN7^T2A>GAL4^; UAS-CSN7::HA* (iv, n= 4) showing rescue of small brain phenotype. Brains are stained with anti-Dlg (green) and anti-Dpn (red) antibodies. A large dashed yellow circle highlights the control brain lobe size, and a smaller dashed red circle indicates the reduced lobe size in mutants (n=number of brains, scale bar= 50µm). v shows the quantification of the relative central brain lobe volume (%). Data is normalized to the *y∗w∗* control. Error bars represent mean ± SD. Statistical significance was determined using a Kruskal-Wallis test followed by Dunn’s multiple comparison test (∗∗p<0.01, ∗∗∗p<0.001, ∗∗∗∗p<0.0001, n.s. = not significant) (C) Mature L3 Larval brains from *y*w*; Insc>GAL4, UAS-mCD8::GFP* control (i, n=21) and *CSN1b* RNAi mediated knock down (RSN2, ii, n= 20 and RSN3, iii, n=27) exhibiting small brain phenotype. Brains are stained with anti-GFP and anti-Dpn antibodies. A large dashed yellow circle highlights the control brain lobe size, and a smaller dashed red circle indicates the reduced lobe size in mutants (n=number of brains, scale bar= 50µm). v shows the quantification of the relative central brain lobe volume (%). Data is normalized to the *y∗w∗* control. Error bars represent mean ± SD. Statistical significance was determined using a Kruskal-Wallis test followed by Dunn’s multiple comparison test (∗∗p<0.01, ∗∗∗p<0.001, ∗∗∗∗p<0.0001, n.s. = not significant).

Next, we analysed whether the small brain phenotype is due to unscheduled apoptosis of NBs and/or premature differentiation of NBs into neurons or glial cells. First, we investigated whether the loss of *CSN7* and *CSN1b* causes a reduction in NB number; therefore, we counted NB number, and interestingly, we found that the number of neuroblasts remained comparable to wild-type (Figure S2-A). Further, to check for an unscheduled apoptosis of NBs, we stained brains with anti-Dcp-1 (*Drosophila* caspase-1). We did not detect any significant increase in Dcp-1-positive NBs in *CSN7* mutants (Controls 0-1 NB and *CSN7* 0-1 NB, Figure S2-B) or in *CSN1b* KD larval brains (Controls 0-1 NB and *CSN1b* 2-4 NB, Figure S2-C). These data suggest that the small brain size of *CSN1b-* and *CSN7-*deficient larvae is not due to apoptosis of NBs. To check whether *CSN1b-* and *CSN7-*deficient NBs lose polarity and prematurely differentiate into neurons or glial cells, we first stained larval brains with Miranda (Mira), a basal polarity marker. We observed that NBs that were undergoing division had a proper basal Mira crescent both in *CSN7* mutants (Figure S3-A) and *CSN1b* KD brains (Figure S3-B), suggesting that *CSN7* and *CSN1b* loss doesn’t affect polarity establishment. Further, we stained the brains with Elav, a pro-neural factor, and Repo, a glial differentiation marker, to check the possibility of premature differentiation of NBs. We did not observe aberrant expression of Elav or Repo in the NBs upon loss of *CSN7* (Elav, Figure S4-A and Repo, Figure S4-B) or *CSN1b* KD (Elav, Figure S5-A and Repo, Figure S5-B), suggesting that *CSN7-* and *CSN1b-*deficient NBs do not undergo premature differentiation. Overall, these data suggest that the small brain phenotype upon the loss of *CSN7* and *CSN1b* is not due to unscheduled apoptosis of NBs and/or premature differentiation of NBs into neurons or glial cells.

### *CSN7* and *CSN1b* are required for growth and proliferation of larval NBs

To further investigate how the loss of *CSN7* and *CSN1b* affects brain development, we examined NBs growth and proliferation. We restricted our study to the central brain NBs as they are the primary drivers of growth in the central brain. We visualised NBs in *CSN7* trans-heterozygotes (*y*w*; CSN7^Mi07174^*/*CSN7^Mi13910^* and *y*w*; CSN7^M13910^*/*CSN7^Mi13910^*/*CSN7 ^T2A>GAL4^*) by staining for Dpn, a NB marker, and Discs-large (Dlg) to mark the cell membrane (Figure 2-A). For *CSN1b* knocked-down brain (*insc-GAL4, UAS-mCD8GFP> UAS-CSN1b Ri*), we used anti-GFP, anti-Dpn and anti-Pros, a neural cell fate determinant (Figure 2-B). We observed that compared to NBs from age-matched control brains, *CSN7-*deficient NBs were significantly smaller in terms of cell diameter (Figure 2-A). We saw a similar reduction in NB size in *CSN1b-*deficient brains (Figure 2-B). These data suggest that the CSN complex is required for appropriate cell growth in larval brain NBs.

**Figure 2:**
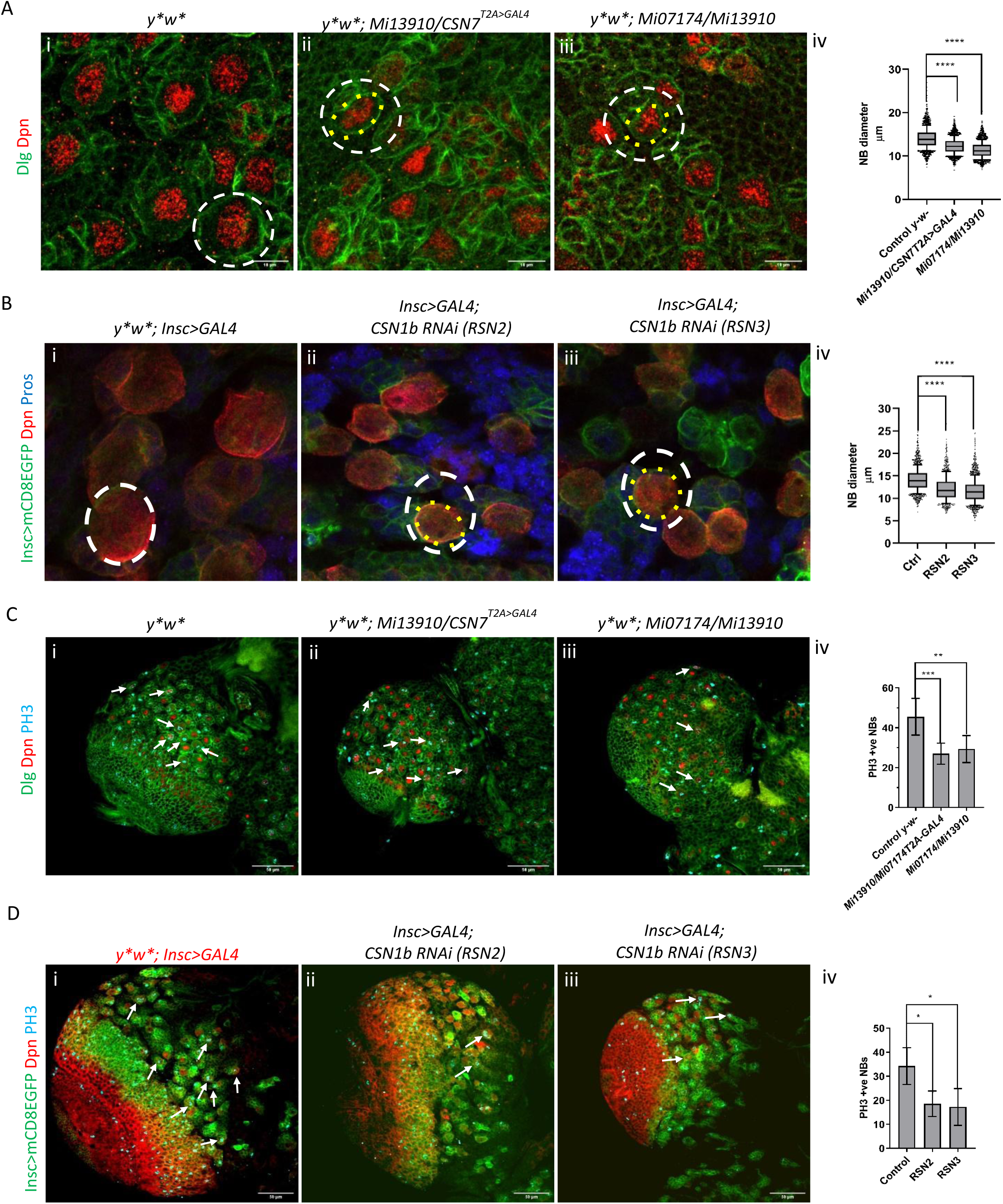
Loss of *CSN7* and *CSN1b* lead to reduced neuroblast size and proliferation. (A) 72 hours ALH larval brains from *y*w\** control (i, n=12), *CSN7* mutants *y*w*; Mi13910/CSN7^T2A>GAL4^* (ii, n=8) and *y*w*; Mi07174/Mi13910* (iii, n=15) stained with anti-Dlg (green) and anti-Dpn (red) antibodies. White and yellow circle mark NBs (scale bar= 10µm). Quantification of NB diameter (μm) is shown in iv. Statistical significance was determined using a Kruskal-Wallis test followed by Dunn’s multiple comparison test (****p < 0.0001) (B) Third-instar larval brains from *y*w*; Insc>GAL4, UAS-mCD8::GFP* control (n=21) and *CSN1b* RNAi mediated knock down (RSN2, n= 22 and RSN3, n= 18) stained with anti-GFP (green), anti-Dpn (red) and anti-Pros (Blue). White and yellow circle mark NBs (scale bar= 10µm). Quantification of NB diameter (μm) is shown in iv. Statistical significance was determined using a Kruskal-Wallis test followed by Dunn’s multiple comparison test (****p < 0.0001). (C) 72 hours ALH larval brains from *y*w\** control (i, n=14), *CSN7* mutants *y*w*; Mi13910/CSN7^T2A>GAL4^* (ii, n=11) and *y*w*; Mi07174/Mi13910* (iii, n=13) stained with anti-Dlg (green), anti-Dpn (red) and anti-PH3 (mitotic marker, blue) antibodies. White arrows mark PH3 positive NBs, mutant brain hemispheres show fewer PH3-positive NBs. Quantification of the average number of PH3-positive NBs per brain lobe is shown in iv. Statistical significance was determined using a Kruskal-Wallis test followed by Dunn’s multiple comparison test (∗∗p<0.01, ∗∗∗p<0.001, ∗∗∗∗p<0.0001) (D) Third-instar larval brains from *y*w*; Insc>GAL4, UAS-mCD8::GFP* control (n=17) and *CSN1b* RNAi mediated knock down (RSN2, n= 16 and RSN3, n= 14) stained with anti-GFP (green), anti-Dpn (red) and anti-PH3 (Blue). White arrows mark PH3 positive NBs, mutant brain hemispheres show fewer PH3-positive NBs. Quantification of the average number of PH3-positive NBs per brain lobe is shown in iv. Statistical significance was determined using a Kruskal-Wallis test followed by Dunn’s multiple comparison test (∗∗p<0.01, ∗∗∗p<0.001, ∗∗∗∗p<0.0001)

**Figure 3:**
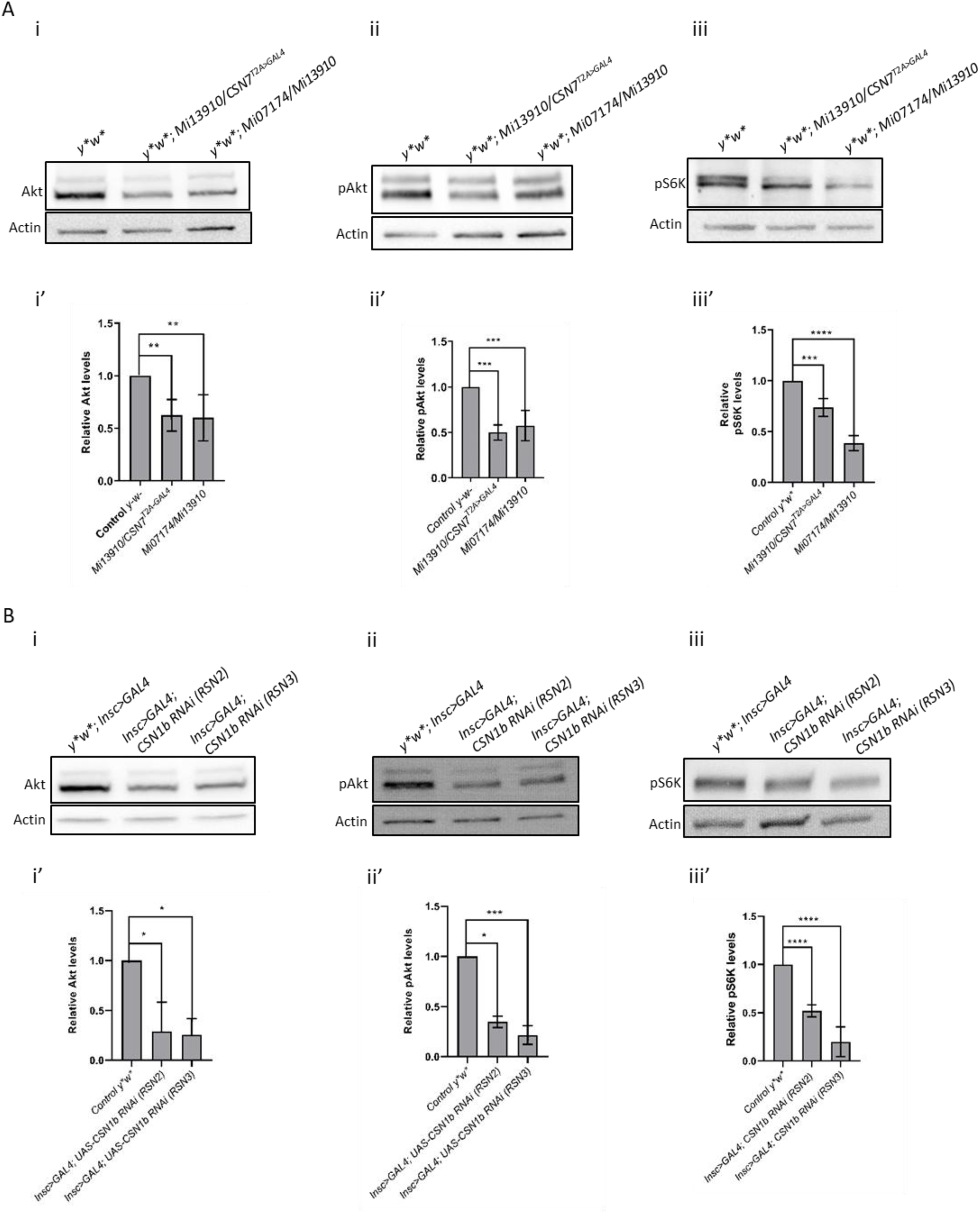
*CSN7* and *CSN1b* deficient NBs have reduced Akt, pAKT and S6k proteins. (A) Western blot analysis of protein extracts from 72 hours ALH larval brains from *y*w\** control, *CSN7* mutants *y*w*; Mi13910/CSN7^T2A>GAL4^* and *y*w*; Mi07174/Mi13910* (n=3). Representative blots are shown for total Akt (i), phosphorylated Akt (pAkt, ii), and phosphorylated S6K (pS6K, iii), Actin was used as a loading control. Accompanying graphs (A-i’, ii’ and iii’) show quantification of relative protein levels normalised to actin and to the control. Consistent significant reductions in Akt, pAkt, and pS6K levels are observed across both *CSN7* mutant heteroallelic combinations compared to control. Statistical significance was determined using a Kruskal-Wallis test followed by Dunn’s multiple comparison test (**p<0.01, ***p<0.001, ****p<0.0001) (B) Western blot analysis of protein extracts from third-instar larval brains from *y*w*; Insc>GAL4, UAS-mCD8::GFP* control and *CSN1b* RNAi mediated knock down (RSN2 and RSN3) (n=3). Representative blots (B-i, ii and iii) show levels of total Akt (i), pAkt (ii) and pS6K (iii). Corresponding quantified relative protein levels are shown in graphs B-i’, ii’ and iii’, normalised to actin and the control. All three signalling markers are significantly reduced in NB-specific knockdown brains. Statistical significance was determined using a Kruskal-Wallis test followed by Dunn’s multiple comparison test (*p<0.05, ***p<0.001, ****p<0.0001).

We next examined whether proliferation of these NBs was also affected upon loss of *CSN7* and *CSN1b.* We stained brains for the mitotic marker PH3 and Dpn, and counted the number of proliferating NBs. We found that compared to control brains, the average number of PH3-positive NBs in *CSN7* trans-heterozygote larval brains were significantly less (Figure 2-C). Similarly, as compared to the control brains, depletion of *CSN1b* in NBs by RNAi also resulted in a reduction in the number of mitotic NBs (Figure 2-D). Since small size and reduced proliferation are characteristics of a quiescent NB, we checked whether *CSN7-* and *CSN1b-* deficient NBs fail to exit the quiescent phase. In *Drosophila,* most NBs become quiescent towards the end of the embryonic phase and re-activate after larval hatching. This re-activation involves cell enlargement and subsequent entry into S-phase followed by mitotic division, with the majority of NBs actively proliferating by 24 hrs ALH (Truman & Bate, 1988; Ito & Hotta, 1992). To test this, we dissected brains from 24 hrs ALH *CSN7* trans-heterozygous and NB-specific *CSN1b* knock-down larvae and stained them for PH3 to count the number of reactivated NBs. We didn’t find any significant differences in the number of PH3-positive NB between control and *CSN7* deficient NBs (Figure S6-A), and control NBs and *CSN1b* deficient NBs (Figure S6-B). These data suggest that *CSN1b-* and *CSN7-*deficient NBs get re-activated after larval hatching at similar time points as controls; therefore, the COP9 signalosome may not be involved in exit from quiescence. However, we cannot rule out the possibility of perdurance of maternally deposited CSN1b/CSN7 proteins that might explain the lack of a phenotype at these early stages of larval development. One possible explanation for the observed phenotype is that once the maternal contribution is depleted, no new protein is translated; *CSN1b-* and *CSN7*-deficient NBs cannot enlarge and therefore cannot enter the cell cycle, resulting in small brains.

Previous studies showed that loss of CSN subunits results in an accumulation of p27 (Bech-Otschir, 2001; Lykke-Andersen *et al*, 2003) and p53 (Lykke-Andersen *et al*, 2003; Tomoda *et al*, 2002) that leads to G1 arrest. Therefore, we decided to check Dmp53 (*Drosophila* p53) and Dacapo (Dap, fly homologue of p27). For Dmp53, we ordered several antibodies, however, none of them gave us a proper signal in immunostaining or on Western blots. Therefore, we checked the expression of Archipelago (Ago), a direct downstream target of p53 that regulates degradation of CycE (Ouyang *et al*, 2011). We performed qRT-PCR on mRNA extracted from appropriate controls and *CSN7*-and *CSN1b*-deficient larval brains. We found that CSN7-deficient brains show a reduced level of Ago mRNA (Figure S6-C). However, there was no significant difference in *Ago* mRNA level in *CSN1b*-depleted brains (Figure S6-D). This data suggest that the small brain phenotype caused by loss of *CSN7* and *CSN1b* may not be due to an accumulation of Dmp53. Next, we checked DAP expression in *CSN7-* and *CSN1b-*deficient brains using an anti-Dap antibody and found no significant change in the number of Dap-positive NBs in either *CSN7* trans-heterozygous mutant brains (Figure S7-A) or in NB-specific *CSN1b* knockdown brains (Figure S7-B), implying that the proliferation defect we observed is not due to the action of Dap.

A previous study in *Drosophila* showed that loss of *CSN7* in the wing disc results in reduced *cycE* expression (Singer *et al*, 2014). To test *cycE* mRNA levels, we performed qRT-PCR on mRNA extracted from controls and *CSN7*- and *CSN1b-*deficient brains. We found a very mild increase in expression levels of *cycE* in *CSN7*-deficient brains compared to controls (Figure S8-A). We didn’t find a significant difference in expression levels of *cycE* between *CSN1b-* deficient brains and controls (Figure S8-B). These data suggest that *CSN7* may have a tissue-specific function, where it regulates *cycE* expression in wing discs but not in developing brains, and the proliferation defects in *CSN7-* and *CSN1b-*deficient NBs may not be due to a decrease in *cycE* expression.

### *CSN7* and *CSN1b* are required for Akt/mTor activity in larval neuroblasts

Our observations so far suggest that larval NBs deficient in *CSN7 and CSN1b* show defects in both proliferation and cell growth. This strongly implicates the Akt/mTOR signalling axis as it is known to be the primary regulator for both cell size and proliferation in various systems (Laplante & Sabatini, 2012; Murakami *et al*, 2004; Yu & Cui, 2016). To investigate this, we quantified relative protein levels of Akt in larval brain extracts from *CSN7* trans-heterozygotes and *CSN1b-*knocked-down brains using Western blotting. We observed a significant reduction in Akt levels in *CSN7* trans-heterozygote brains as compared to controls (Figure 4-A, i and i’). Similarly, extracts from *CSN1b* knocked-down brains also showed a significant decrease in Akt (Figure 4-B, i and i’). Since Akt levels were reduced in both *CSN1b-* and *CSN7-*deficient brains, we checked whether it is regulated at the transcriptional level. For this, we performed qRT-PCR on mRNA extracted from controls, *CSN7* trans-heterozygotes and *CSN1b* knocked-down brains. We didn’t find any significant difference in expression levels of *akt* between *CSN1b-* and *CSN7-*deficient brains and respective controls (Figure S9). These data suggest Akt is regulated at the post-translational level.

**Figure 4:**
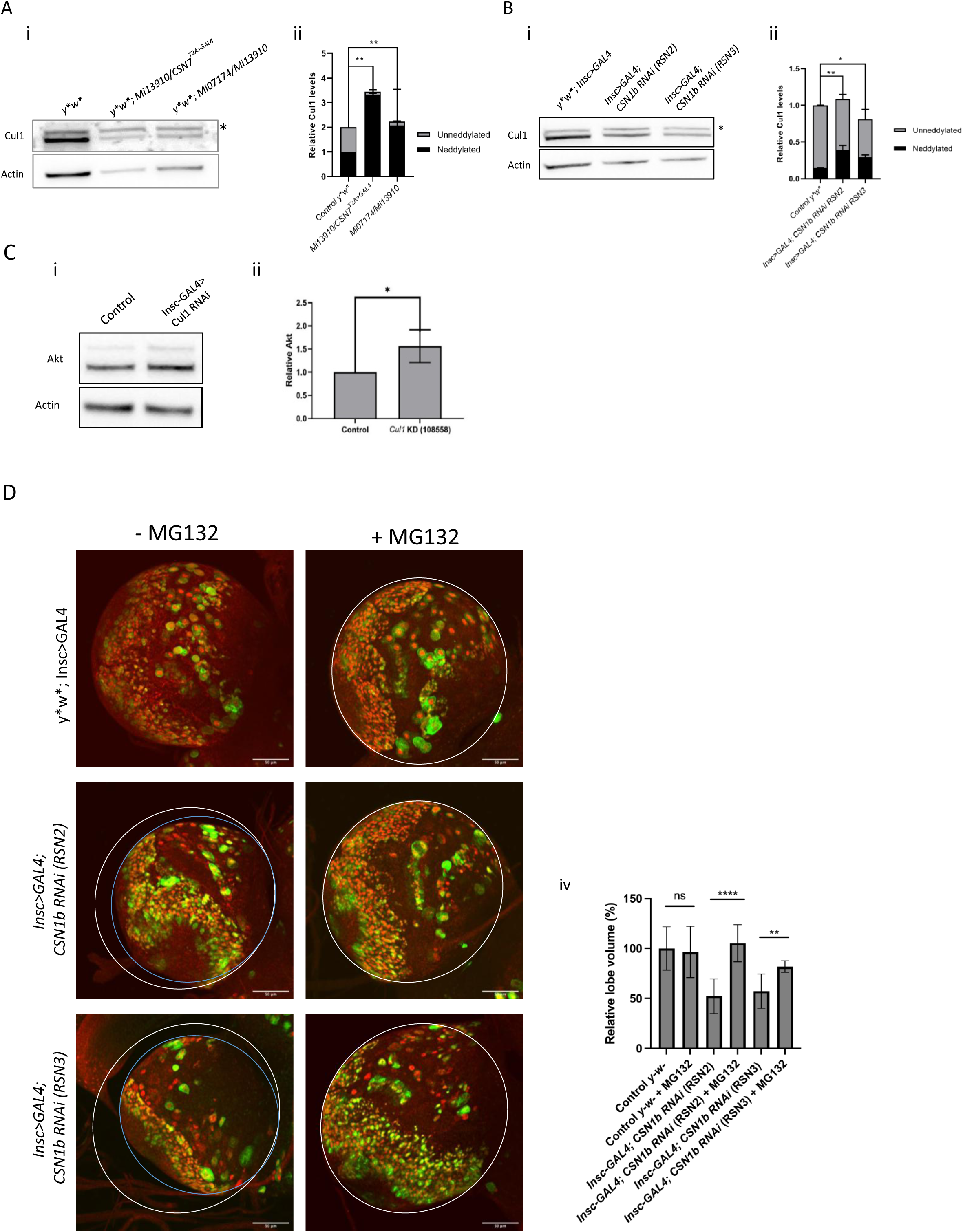
*CSN7* and *CSN1b* deficient brains accumulate neddylated Cul1. (A) Western blot analysis of protein extracts from 72 hours ALH larval brains from *y*w\** control, *CSN7* mutants *y*w*; Mi13910/CSN7^T2A>GAL4^* and *y*w*; Mi07174/Mi13910* (n=3). Blot probed with anti-Cul1 antibody (i) shows increased intensity of the top Cul1 band (marked by asterisk), corresponding to neddylated Cul1 in mutant brain extracts. Actin was used as a loading control. Quantification of relative levels of neddylated (black portion of bar) and unneddylated (gray portion of bar) Cul1is shown in ii (B) Western blot analysis of protein extracts from third-instar larval brains from *y*w*; Insc>GAL4, UAS-mCD8::GFP* control and *CSN1b* RNAi mediated knock down (RSN2 and RSN3) (n=3). Blot was probed with anti-Cul1 and anti-Actin antibodies. An accumulation of the higher molecular weight, neddylated Cul1 (asterisk) is observed in knockdown conditions compared to control (i). Quantification of relative levels of neddylated (black portion of bar) and unneddylated (gray portion of bar) Cul1is shown in ii. All data are presented as mean ± SD. Statistical significance was determined using a Kruskal-Wallis test followed by Dunn’s multiple comparison test ((*p<0.05, **p<0.01) (C) Western blot of extracts from age matched brains from control (*Insc>GAL4*) and NB-specific *Cul1* knock down larvae. Actin was used as a loading control. Quantification of relative Akt protein levels normalised to actin. Knockdown of Cul1 leads to a significant increase in total Akt levels in the larval brain. Data are presented as mean ± SD. Statistical significance was determined using a two-tailed Mann-Whitney U test (*p<0.05). (D) Third-instar larval brains from *UAS-Dicer; Insc>GAL4, UAS-mCD8::GFP* control (i, n=5), *UAS-Dicer; Insc>GAL4, UAS-mCD8::GFP*; RSN2 (ii, n=5), *UAS-Dicer; Insc>GAL4, UAS-mCD8::GFP*; RSN3 (iii, n=5) and MG132 fed *UAS-Dicer; Insc>GAL4, UAS-mCD8::GFP* control (i’, n=5), *UAS-Dicer; Insc>GAL4, UAS-mCD8::GFP*; RSN2 (ii’, n=5), *UAS-Dicer; Insc>GAL4, UAS-mCD8::GFP*; RSN3 (iii’, n=5) stained with anti-Dpn (red), anti-Mira (green) and anti-PH3 (blue) antibodies. White circle shows the control brain size and blue circles in ii and iii highlights the small brains. (scale bar= 50µm). (iv) shows the quantification of the relative central brain lobe volume (%). Data is normalized to the *y∗w∗* control. Error bars represent mean ± SD. Statistical significance was determined using a Kruskal-Wallis test followed by Dunn’s multiple comparison test (∗∗p<0.01, ∗∗∗p<0.001, ∗∗∗∗p<0.0001, n.s. = not significant).

Further, to determine whether this reduction in total Akt translates to a functional deficit in pathway activity, we similarly assessed the levels of pAkt, the active form of the kinase. Consistent with our results for Akt, we observed that pAkt levels were also significantly diminished in extracts from both *CSN7-*deficient brains (Figure 4-A, ii and ii’) and *CSN1b-* deficient brains (Figure 4-B, ii and ii’). To confirm whether this deficit in Akt signalling extends to the downstream mTOR pathway as well, we quantified relative levels of pS6K. We found that pS6K levels were significantly decreased in the larval brain extracts from both *CSN7* trans-heterozygotes (Figure 4-A, iii and iii’) and *CSN1b* knocked-down brains (Figure 4-B, iii and iii’), suggesting severe attenuation of the mTOR pathway in these animals. Overall, our data suggest that loss of *CSN7/CSN1b* regulates Akt/mTor signalling in NBs by regulating Akt stability.

### CSN7 and CSN1b stabilise Akt levels via neddylation of Cul1

The CSN complex is a known regulator of the ubiquitin-proteasome system by deneddylating Cullin-RING-ligases (CRLs) (Cope *et al*, 2002; Wei & Deng, 2003). A previous report in *Drosophila* showed that Cul1 ligase facilitates ddaC neuronal dendrite pruning by regulating InR/PI3K/TOR pathway via ubiquitination and degradation of Akt (Wong *et al*, 2013). We hypothesised that the CSN complex might regulate Akt/mTOR mediated larval NBs growth by inactivating Cul1 through deneddylation, which in turn stabilizes Akt. To test this, we performed Western blotting to examine the neddylation status of Cul1 in larval brain extracts using anti-Cul1 antibody that recognises both neddylated and deneddylated forms. In control brains, Cul1 is present in both the neddylated (active) and un-neddylated (inactive) forms (Figure 4-A, i and ii). However, in larval brain extracts from *CSN7* trans-heterozygotes, we observed a dramatic accumulation of the neddylated variant (NEDD8-Cul1) (Figure 4-A, i and ii). A similar though less pronounced significant increase in NEDD8-Cul1 was observed in extracts from *CSN1b* knocked-down brains (Figure 4-B, i and ii). This accumulation of neddylated Cul1 suggests that the CSN complex is required for Cul1 dennedylation and that its loss results in the accumulation of potentially hyper-active Cul1 that may be leading to rapid Akt turnover. To further check whether Cul1 negatively regulates Akt in the larval brain, we performed an NB-specific knockdown of *Cul1* using *insc-GAL4, UAS-mCD8GFP* and *UAS-Cul1* RNAi (v108558) in a wild-type background and assessed Akt levels in larval brain extracts. We observed a significant increase in total Akt protein levels in *Cul1-*deficient larval brains suggesting that Cul1 negatively affects Akt levels (Figure 4-C). Next, we checked whether the small brain phenotype upon *CSN1b* knockdown in NBs is due to proteasome-mediated degradation of Akt. For this, we fed MG132, an inhibitor of 26S proteasome that inhibits the degradation of ubiquitin-conjugated proteins, to larvae and assessed brain size and found that the brain size was rescued (Figure 4-D). Taken together, these data suggest that the accumulation of neddylated Cul1 in *CSN1b-*deficient larval brains leads to excessive E3 ligase activity, leading to degradation of Akt that eventually suppresses the Akt-TOR pathway in NBs.

For further validation of the hypothesis that CSN regulates Akt-TOR pathway via Cul1 in NB, we ectopically expressed Akt (*insc-GAL4, UAS-mCD8GFP> UAS-Akt; UAS-CSN1b Ri*) in *a CSN1b-*knockdown background and found that overexpression of Akt rescues the small brain and NB size phenotypes (Figure 5A-i, ii, iv and v). Next, we knocked down *Cul1* in *CSN1b-* deficient NBs (*insc-GAL4, UAS-mCD8GFP> UAS-cul1 Ri; UAS-CSN1b Ri*) and found that it also rescues NB size and small brain phenotype (Figure 5A-iii and v). Taken together, our data suggest that the CSN complex regulates NBs size and proliferation by modulating Akt levels by deactivating Cul1 through deneddylation (Figure 5-B).

**Figure 5:**
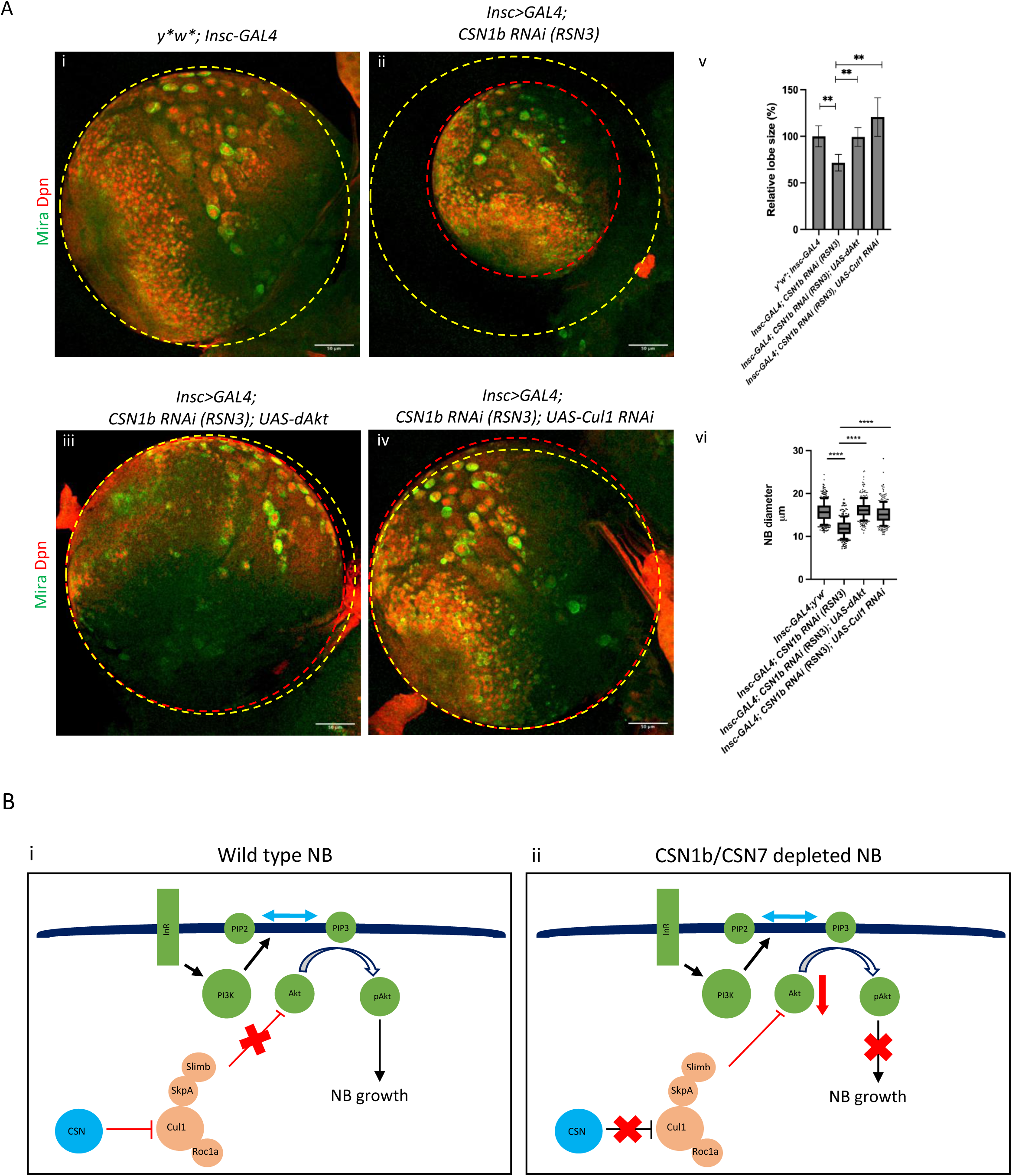
Overexpression of Akt and downregulation of Cul1 rescue small brain phenotype. (A) Third-instar larval brains from *UAS-Dicer; Insc>GAL4, UAS-mCD8::GFP* control (i, n=5), *UAS-Dicer; Insc>GAL4, UAS-mCD8::GFP*; RSN3 (ii, n=5), *UAS-Dicer; Insc>GAL4/ UAS-Cul1*; RSN3 (iii, n=7); and *UAS-Dicer; Insc>GAL4/ UAS-Akt Ri*; RSN3 (iv, n=6). Brains are stained with anti-Mira and anti-Dpn antibodies (scale bar= 50µm) (v) shows the quantification of the relative central brain lobe volume (%). Data is normalized to the control (vi) shows the quantification of the relative NB diameter. Error bars represent mean ± SD. Statistical significance was determined using a Kruskal-Wallis test followed by Dunn’s multiple comparison test (∗∗p<0.01, ∗∗∗p<0.001, ∗∗∗∗p<0.0001, n.s. = not significant) (B) Schematic diagram showing regulation of Akt levels in control NB (i) and *CSN1b/CSN7* depleted NB (ii). In control NB (i), Cul1 is deneddylated and inactivated by CSN. As a result, Akt levels are maintained, which results in helps in NB growth (ii) shows the senerio where in CSN1b or CSN7 depleted NBs, Cul1 is no longer inactivates by CSN. This in turn leads to rapid degradation of Akt, which results in small size NBs and brains.

## Discussion

There is considerable literature demonstrating the importance of the CSN complex in various developmental processes in *Drosophila*, including wing disc development (Suisse *et al*, 2017; Doronkin *et al*, 2003), stem cell differentiation (Qian *et al*, 2015), oogenesis (Doronkin *et al*, 2002), and neural competence (Huang *et al*, 2014), however, there have been no studies describing any role for the CSN in neurogenesis. In this study, we establish a role for the CSN complex in brain development in *Drosophila*. We show that loss of CSN subunits, CSN7 and CSN1b, results in a severe reduction in brain volume and NB growth (Figure 1). Our investigations show that this small brain phenotype arises from cell-autonomous defects in NB growth and proliferation (Figure 2), and are not due to increased apoptosis (Figure S2-B and C), polarity defects (Figure S3-B and C) or premature differentiation into neurons (Figure S4-A and B) or glia (Figure S5-A and B), or a failure to exit from quiescence (Figure S6 A and B). Additionally, we also show that the phenotype does not arise from elevated levels of p53 (via Archipelago) (Figure S6-A and D) or Dacapo (Figure S7).

Re-growth of NBs post-division is a necessary prerequisite to enter into successive divisions (Betschinger *et al*, 2006; Hartenstein *et al*, 1987) during larval neurogenesis. The Hippo pathway and the Akt/mTor pathway are the main drivers of cell size during development (Laplante & Sabatini, 2012; Murakami *et al*, 2004; Yu & Cui, 2016). The Hippo pathway is necessary for exit of NBs from quiescence (Ly *et al*, 2019). We found that *CSN7-* and *CSN1b-* deficient NBs successfully exit quiescence and get reactivated after larval hatching at similar time points as controls, suggesting that CSN1b and CSN7 may not be involved in exit from quiescence; therefore, the Akt/mTor pathway becomes strongly implicated. However, we cannot rule out the possibility of perdurance of maternally deposited CSN1b/CSN7 that might explain the lack of a phenotype at these early stages of larval development. One explanation for the observed phenotype could be that once the maternal contribution is depleted and no new protein is translated, *CSN1b-* and *CSN7-*deficient NBs cannot enlarge and therefore cannot enter the cell cycle, resulting in small brains.

Therefore, we assessed Akt and the activated form pAkt protein levels and found that both were markedly reduced in the developing larval brain upon loss of either *CSN7* or *CSN1b* (Figure 3). We also observed a corresponding reduction in mTOR activity as indicated by decreased pS6K levels (Figure 3). Crucially, transcript levels of Akt remain unchanged upon loss of CSN7/CSN1b, indicating that the CSN complex acts post-transcriptionally to maintain Akt stability (Figure S9). Further, targeted ectopic expression of Akt in the developing larval brain in *a CSN7-* or *CSN1b-*deficient background restores brain lobe volume (Figure 5), demonstrating that attenuation of the Akt-mTOR pathway is the major driver of the phenotype, if not exclusive. While cell growth and cell proliferation are often conflated as being the same phenomenon, it is important to note that they are distinct processes (Kaldis, 2016). In the case of larval neurogenesis, cell growth must necessarily precede cell division in NBs. This is consistent with our data wherein we observe smaller NBs that fail to proliferate, suggesting that the failure to grow in size prevents or inhibits proliferation.

The CSN complex is a well-known deneddylase for Cullin-RING ligases and is highly conserved (Cope *et al*, 2002; Wei & Deng, 2003). We therefore asked whether its loss in NBs of the developing larval brain might perturb CRL function in a manner that would impact the Akt/mTor axis. We observed that depletion of either *CSN7* or *CSN1b* in larval NBs results in a pronounced accumulation of neddylated Cul1, which is the active form (Figure 4-A and B). Accumulation of neddylated Cullins is a clear biochemical indicator of hyperactivity. Data from our study support a model wherein a hyperactive Cul1-SCF is responsible for excessive Akt turnover. First, NB-specific knockdown of *Cul1* enhances Akt levels in the developing larval brain in an otherwise wild-type background (Figure 4-D). Second, feeding MG132 to larvae with NB-specific *CSN1b* knockdown rescues the small brain phenotype (Figure 4-C). A previous report has shown that Cul1 regulates Akt levels during neuronal pruning in early pupae (Wong *et al*, 2013). Our findings suggest an analogous mechanism in NBs for regulating Akt levels during larval neurogenesis at an earlier stage of development. The fact that both CSN7 and CSN1b are essential for brain development suggests that the entire holocomplex is essential for Akt regulation in NBs.

While the CSN complex has been studied extensively in the context of protein turnover in general (Von Arnim, 2003) as well as cell-cycle control (Zhang *et al*, 2008; Lykke-Andersen *et al*, 2003), its specific function in sustaining a growth-promoting stimulus during larval neurogenesis has not been reported so far. Our work demonstrated a regulatory role for the CSN complex during larval neurodevelopment where it restrains Cul1-SCF activity via deneddylation, thereby allowing the Akt/mTOR pathway to promote growth of NBs and subsequent proliferation (Figure 5-B). This regulatory function is crucial for Akt/mTOR activity in larval NBs and therefore indispensable for neurodevelopment in *Drosophila*. Our study has not tested whether the mechanism presented herein is conserved in vertebrate or mammalian models. However, given the highly conserved nature of both the CSN complex and the Akt/mTor pathway, there is a good degree of probability that the basic mechanism may be conserved, albeit with varying degrees of complexity.

## Materials and methods

### Fly stocks and husbandry

All flies were reared at 25°C unless otherwise specified on standard media. For knockdown experiments, flies were grown at 29°C. Where required, flies were grown in cages and 10mm dishes containing standard media were used for timed collection of embryos.

For dissection of larval brains, wandering L3 larvae were staged visually and dissected. When larvae of very specific age were required (± 3 hours), plates with timed embryo collections were used.

Genotyping of larvae with GFP balancers was performed using an Olympus stereo microscope where required.

The following table lists the fly strains used in this study

| Genotype | Source |
| --- | --- |
| *y[1] w[\*]* | Bellen lab |
| *UAS-Dicer; insc-GAL4>CD8-GFP* | Neumuller et al., 2011 |
| *y[1] w[\*]; P{w[+mC]=Act5C-GAL4}17bFO1/TM6B, Tb[1]* | BDSC |
| *y[1] w[\*]; P{w[+mC]=Tub-GAL4}/TM6B, Tb[1]* | BDSC |
| *w1118; P{GD11099}v34727* | VDRC |
| *y1 v1; P{TRiP.JF02612}attP2* | BDSC |
| *y[1] v[1]; P{y[+t7.7] v[+t1.8]=TRiP.HM05119}attP2* | BDSC |
| *y[1] sc[\*] v[1]; P{y[+t7.7] v[+t1.8]=TRiP.HMS00242}attP2* | BDSC |
| *y[1] v[1]; P{y[+t7.7] v[+t1.8]=TRiP.GL01169}attP2* | BDSC |
| *y[1] v[1]; P{y[+t7.7] v[+t1.8]=TRiP.JF03159}attP2* | BDSC |
| *y[1] v[1]; P{y[+t7.7] v[+t1.8]=TRiP.HMJ30047}attP40* | BDSC |
| *y[1] v[1]; P{y[+t7.7] v[+t1.8]=TRiP.GL01151}attP2* | BDSC |
| *y[1] v[1]; P{y[+t7.7] v[+t1.8]=TRiP.HMS02392}attP40* | BDSC |
| *y[1] sc[\*] v[1] sev[21]; P{y[+t7.7] v[+t1.8]=TRiP.GL00492}attP2* | BDSC |
| *w1118; P{GD12914}v40690* | VDRC |
| *w1118; P{GD12914}v40691* | VDRC |
| *y[1] sc[\*] v[1] sev[21]; P{y[+t7.7] v[+t1.8]=TRiP.HMS00073}attP2* | BDSC |
| *y[1] sc[\*] v[1] sev[21]; P{y[+t7.7] v[+t1.8]=TRiP.HMS00243}attP2/TM3, Sb[1]* | BDSC |
| *y[1]w[\*]; Mi{y[+mDint2]=MIC}CSN7[MI07174]/Cyo, Ubi-GFP* | BDSC |
| *y[1] w[\*]; Mi{y[+mDint2]=MIC}CSN7[MI13910]/Cyo, Ubi-GFP* | BDSC |
| *y[1] w[\*]; Mi{Trojan-GAL4.1}CSN7[MI07174-TG4.1]/Cyo, Ubi-GFP* | BDSC |
| *w[1118]; P{w[+mC]=UAS-S6k.M}2/Cyo* | BDSC |
| *UAS-dAkt/Cyo* | BDSC |
| *y[1] w[\*]; P{w[+mC]=UAS-CSN7::HA}3* | This study |
| *Y[1] w[\*]; Mi{PT-GFSTF.0}CSN7<sup>MI7174-GFSTF.0</sup>/SM6a* | This study |

### Transgenic line generation

A full-length CSN7-HA cDNA fragment was commercially synthesized (Genscript). Fragment: CSN7-HA was digested and cloned into the pUASTattB vector between EcoRI and XbaI sites. To generate transgenic lines, pUAST-CSN7::HA was injected into *y^1^w^∗^*; *VK33* embryos. Injected adults were screened for transgenes with red eyes.

### Immunostaining and Western Blotting

Larval brains were dissected in Phosphate Buffered Saline (PBS) and then fixed in 4% Paraformaldehyde (HiMedia TCL119) for 20 minutes; samples were then rinsed thrice in PBST (PBS + 0.3% TritonX) at room temperature and then blocked using 10% Normal Goat Serum (NGS) for 1 hour. Incubation with the primary antibodies was done overnight at 4°C, followed by three rinses with PBST at room temperature. Brains were afterward incubated with the secondary antibodies at room temperature for 2 hours, rinsed thrice with PBST, and then mounted in ProLongTM Gold Antifade mounting media (Invitrogen, P36934) on glass slides and sealed with a coverslip for microscopy. For Westerns, larval brains were dissected and isolated in PBS and homogenised in lysis buffer (50mM Tris, 100mM NaCl, 5mM EDTA, 1x PIC) and then boiled with 1x Laemmeli buffer with dye. Samples were separated on SDS-PAGE and transferred onto a nitrocellulose membrane using a semi-dry transfer (Trans-blot turbo transfer system, BioRad 1704150). Blots were subjected to blocking for one hour at room temperature with either 5% BSA (Himedia, MB083) or 5% Blotto (Santacruz, sc2325) and then probed with the requisite primary antibody overnight at 4°C. Blots were then washed with Tris Buffered Saline with Tween20 (TBST, 0.1% Tween20) thrice, probed with the appropriate secondary antibody for 2 hours at room temperature and washed thrice again with TBST. Blots were developed using SuperSignal WEST Pico plus (ThermoFischer, 34579) and visualised on a chemidoc (Fusion eVo-6, Vilber GmbH). Densitometric measurements were performed using Fiji.

### BrdU incorporation

Larvae of appropriate age were grown on media containing 20 μg/mL of BrdU (ThermoFischer B23151) for 2 hours and then dissected in 1x PBS. The brains were processed for immunostaining using anti-BrdU antibody as described previously with the addition of a 30 minute treatment with 2.2N HCl in 1x PBS-TritonX followed by three washes with 100mM Borax buffer before blocking.

### Image acquisition and brain volume analysis

Larval brains were imaged using confocal microscopy either on a Zeiss LSM 880 or a Leica SP8. Laser power, gain etc. were calibrated at the start of an experiment and maintained constant throughout. Complete brain lobes were imaged using a 40x oil immersion lens (Plan-Apochromat 40x/1.4 Oil DIC M27) and when necessary, a 60x oil immersion lens (Plan-Apochromat 63x/1.4 Oil DIC M27) was used. Quantification was performed on Fiji or by 3D reconstruction using Imaris (Bitplane). All quantifications were performed on images wherein pixel values remained unaltered. For quantification of NB numbers, image stacks of larval brain lobes were manually segmented and NBs were marked using the Region of Interest (ROI) function. Feret’s diameter measurements were used for NB size and average mean grey values for segmented NBs were used where fluorescence intensity was needed. For measurements of brain lobe volume, brain lobes were manually segmented using a 3D reconstruction on Imaris. The entire brain lobe was imaged as a stack of 1μm slices. The brain lobes were manually segmented on each plane and subsequently rendered into a single volumetric object. Using the voxel measurement data from the raw image, the volume of the reconstructed 3D brain lobe was computed. A volumetric object was generated using the segmented image which was then used to derive the volume of the brain lobes in µm^3^.

### RNA isolation and quantitative PCR

Larvae were sorted and rinsed in DEPC-treated MQ water, and brains were dissected in DEPC-treated PBS. Brains were homogenised in TRIzol (Ambion, Cat. No. 15596018), followed by chloroform extraction (HiMedia, Cat. No. MB109) and centrifugation at 15,000 rpm for 5 min at 4 °C to achieve phase separation. The aqueous phase was transferred to a fresh tube, and RNA was precipitated using isopropanol (HiMedia, Cat. No. MB063). The RNA pellet was washed with 70% ethanol, air-dried, and resuspended in DEPC-treated water (MB076-HiMedia).RNA concentration was determined, and 1 µg RNA was treated with DNase I (Invitrogen, Cat. No. 18068-015) in a 10 µL reaction following the manufacturer’s instructions. Samples were incubated at room temperature for 15 min, and the reaction was terminated by adding EDTA and incubating at 65 °C for 10 min. First-strand cDNA synthesis was performed using the High-Capacity cDNA Reverse Transcription Kit (Thermo Fisher Scientific, Cat. No. 4368814) with random primers in 20 µL reactions. Reverse transcription was carried out at 25 °C for 10 min, 37 °C for 120 min, and 85 °C for 5 min. No reverse transcriptase controls were included to confirm the absence of genomic DNA contamination. Synthesised cDNA was stored at −20 °C until further use. Quantitative PCR was performed using iTaq Universal SYBR Green SuperMix (Bio-Rad, Cat. No. 1725124) in 20 µL reactions containing 1 µL of 1:16 diluted cDNA and gene-specific primers. Amplification was conducted with an initial denaturation at 95 °C for 3 min, followed by 40 cycles of 95 °C for 10 s and 60 °C for 30 s.

### Statistical analyses

For the analysis, non-parametric tests were employed as the datasets did not follow a normal distribution. For experiments involving three genotypes (control and two trans-heterozygous combinations or control and two different RNAi lines), a Kruskal-Wallis test was used followed by Dunn’s multiple comparison test to identify statistical differences between the experimental groups and the control. For experiments involving only two genotypes a two-tailed Mann-Whitney U test was performed. Statistical significance was set at p < 0.05. Specific sample sizes (n) and statistical tests for each experiment are detailed in the corresponding figure legends.

### Antibodies

| Antibodies | Source | Application |
| --- | --- | --- |
| Mouse aPKC | Sanatacruz sc17781 | IF |
| Rabbit Actin | CST4967S | WB |
| Rabbit Akt | CST 9272Ss | WB |
| Rabbit pAkt | CST 4060 | WB |
| Mouse BrdU | DSHB G2G4 2314035 | IF |
| Rabbit DCP-1 | CST | IF |
| Rabbit Cullin 1 | Invitrogen 718700 | WB |
| Rabbit Dacapo | C. Lehner | IF |
| Mouse Dlg | DSHB | IF |
| Rat Deadpan | Abcam 95173 | IF |
| Guinea pig Deadpan | H.Wang | IF |
| Rat Elav | DSHB 528218 | IF |
| Chicken GFP | Invitrogen A10262 | IF/WB |
| Mouse HA | CST 2367s | WB |
| Rabbit Phospho-Histone H3 (Ser10) | Upstate32219 | IF |
| Mouse mCherry | Invitrogen M11217 | IF |
| Rat Miranda | Abcam ab197788 | IF |
| Mouse Miranda | H Wang | IF |
| Rabbit PKCzeta | Abcamab59384 | IF |
| Mouse Prospero | DSHBmr1a s | IF |
| Rabbit phospho- S6K | CST9209S | WB |

## Author contributions

N.J., P.P. and S.N-J designed the study; N.J., P.P., S.A., D.B. and S.N-J performed the experiments; N.J., P.P., S.A., D.B., M.J. and S.N-J analyzed the data; and N.J., P.P., S.A., D.B., M.J. and S.N-J wrote the manuscript.

## Acknowledgments

We thank Hugo J Bellen for various fly lines used in this study. We are grateful to Hongyan Wang and Christian Lehner for sharing antibodies. We thank Bloomington *Drosophila* Stock Center and Vienna *Drosophila* Stock Center for stocks. We thank Dr. Deepti Trivadi, *Drosophila* injection facility at Bangalore LifeScience Cluster, NCBS-TIFR for embryo injections. We thank Reshmi Verghese, Brinda Palliyana and Shalaka More for technical help. We thank Bharathi, Srikath, Krishna, and Rani from the fly facility, and Subbalakshmi, Suman, and Mahesh from the microscopy facility. SN-J started the protein expression screen in the lab of Hugo J Bellen, and we thank him for his generous support. We thank SN-J lab for fruitful discussions. NJ is supported by CSIR-SRF, CRG-SERB (CRG/2022/007401), and CSIR-CCMB (IHP240003), PP is supported by DBT/Wellcome India alliance (IA/I/18/1/503629) and CSIR-SRF. DB and MJ are supported by the Department of Atomic Energy, Government of India (Project Identification No. RTI4007). SN-J is supported by DBT/Wellcome India alliance (IA/I/18/1/503629), CRG-SERB (CRG/2022/007401) and CSIR-CCMB grant (IHP240003).

