## Supplemental Figures for "COP9 complex maintains neuroblast growth and proliferation by regulating Akt/mTor pathway"

### Figure S1

A

| S. no. | Gene | Flybase ID | Human orthologue |
|---|---|---|---|
| 1 | Roc2 | FBgn0044020 | RNF7 |
| 2 | Ac78c | FBgn0024150 | ADCY8 |
| 3 | Jumu | FBgn0015396 | FOXN4 |
| 4 | bon | FBgn0023097 | TRIM24 |
| 5 | bs | FBgn0004101 | SRF |
| 6 | ed | FBgn0000547 | NPHS1 |
| 7 | fipi | FBgn0031627A | GLON5 |
| 8 | kst | FBgn0004167 | SPTBN5 |
| 9 | CSN7 | FBgn0028836 | COPS7A/COPS7B |
| 10 | Timeout | FBgn0038118 | TIMELESS |
| 11 | kat80 | FBgn0040207 | KATNB1 |
| 12 | Atpα | FBgn0002921 | ATP1A3 |
| 13 | wrd | FBgn0042693 | PPP2R5D |
| 14 | Hrb98DE | FBgn0001215 | HNRNPA1L2 |
| 15 | CSN7 | FBgn0028836 | COPS7B |
| 16 | Cdep | FBgn0265082 | FARP2 |
| 17 | CG42321/ATP8A | FBgn0259221 | ATP8A1 |
| 18 | Rm62 | FBgn0003261 | DDX17 |
| 19 | Cdi | FBgn0004876 | TESK2 |
| 20 | Eip75B | FBgn0000568 | NR1D2 |
| 21 | pyd | FBgn0262614 | TJP1 |
| 22 | Eip93F | FBgn0264490 | LCORL |
| 23 | Alg-2 | FBgn0086378 | PDCD6 |
| 24 | UbcE2M | FBgn0035853 | UBE2M |
| 25 | ctrip | FBgn0260794 | TRIP12 |
| 26 | CG7337/Wdr62 | FBgn0031374 | MAPKBP1/WDR62 |
| 27 | CG34401/Dora | FBgn0085430 | ZSWIM8 |
| 28 | sba | FBgn0016754 | MBD5 |
| 29 | vib | FBgn0267975 | PITPNB |
| 30 | siz | FBgn0026179 | QSEC1 |

B

*Gene::GFP::Gene* (X)*; UAS-DegradFP/Insc-GAL4*
*Gene::GFP::Gene* (II)*, UAS-DegradFP/Wor-GAL4*
*Insc-GAL4/UAS-DegradFP; Gene::GFP::Gene (III)*

↓ 29˚C

Lethal

↓

Brain size analysis

C

| S. no. | Gene | Lethal/viable | Brain size |
|---|---|---|---|
| 1 | Roc2 | lethal | small |
| 2 | kat80 | lethal | small |
| 3 | vib | lethal | small |
| 4 | Cdi | lethal | small |
| 5 | Cdep | lethal | small |
| 6 | CSN7 | lethal | small |
| 7 | CG3773/Wdr62 | lethal | small |
| 8 | Hrb98DE | lethal | small |
| 9 | Atpα | lethal | small |
| 10 | CG42321/ATP8A | lethal | small |
| 11 | sba | lethal | small |
| 12 | wrd | lethal | small |
| 13 | CG34401/Dora | lethal | small |

D

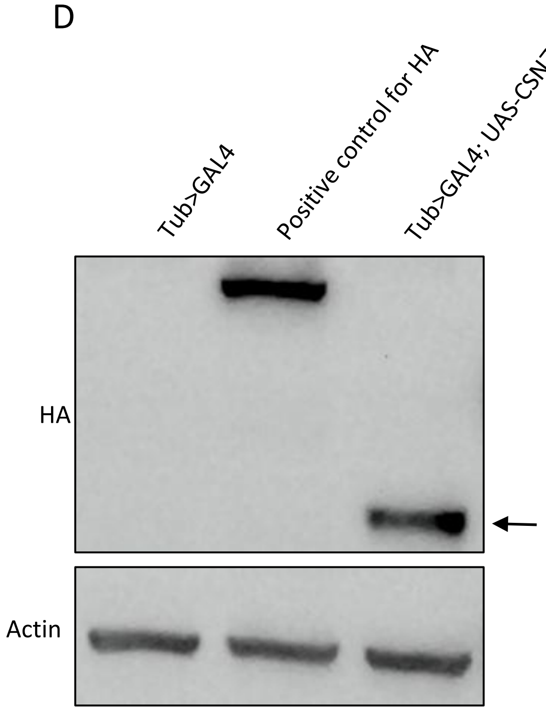

**Figure S1: An expression based screen to identify genes critical for brain development in *Drosophila*.** (A) Table listing genes from expression based screen with expression in developing larval brain (B) Schematic of the gene knockdown and screening assay, to specifically knock down the endogenous protein of interest in larval neuroblasts, *Insc-GAL4* or *Wor-GAL4* was used to drive *UAS-degradFP* in a background homozygous for the MiMIC-protein-trap line for genes listed in (A), (C) a curated list of candidate genes identified from the screen. For each gene, the phenotype observed upon specific protein knockdown in larval neuroblasts is listed (D) whole larval extracts from Tub>GAL4, flies expressing HA tagged protein (positive control) and *Tub>GAL4; UAS-CSN7-HA*. Blots are probed with anti-HA and anti-Tubulin (loading control) antibodies.

**Figure S2**

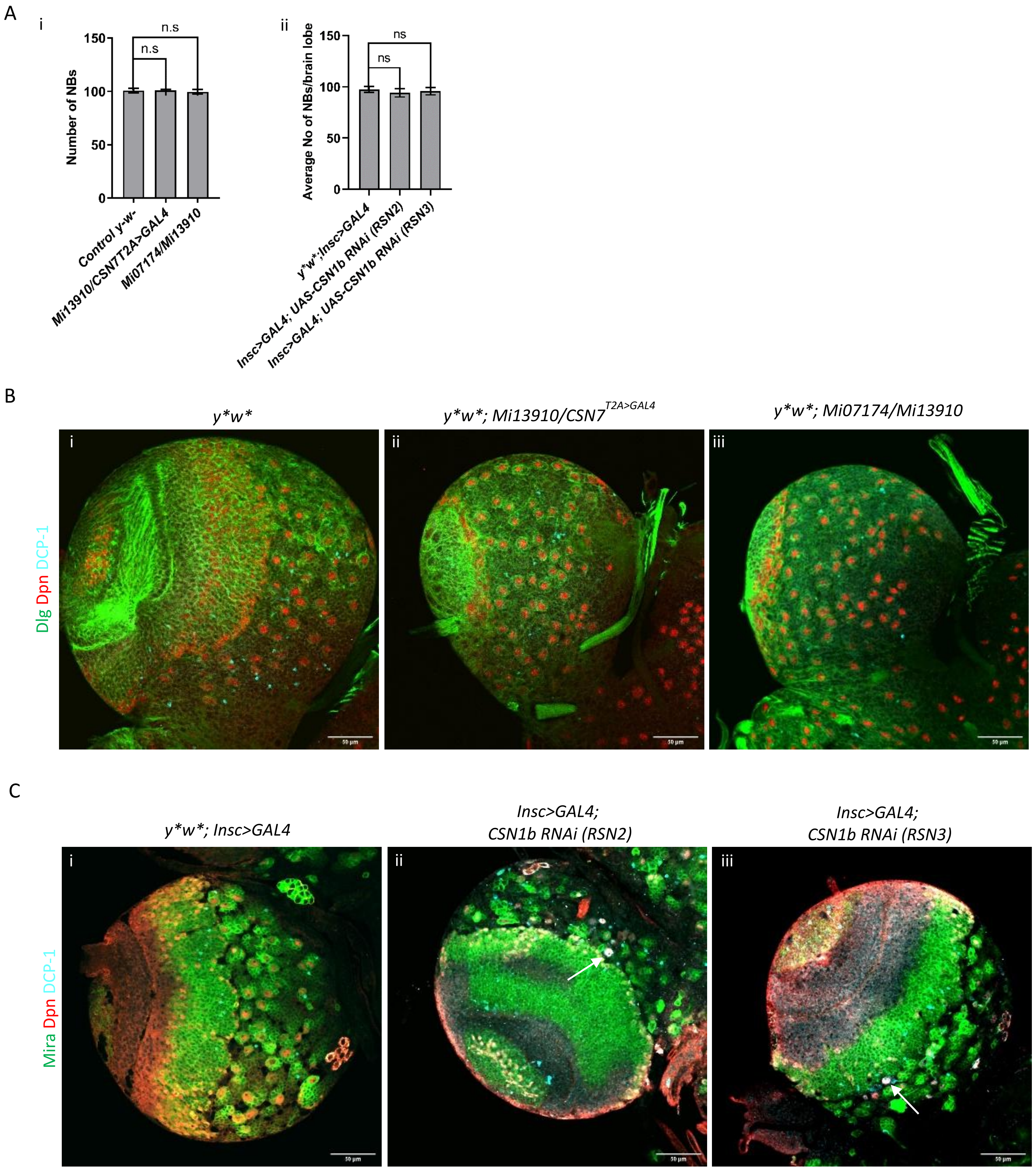
A
i
n.s
n.s
Number of NBs
150
100
50
0
Control y-w-
Mi13910/CSN7T2A>GAL4
Mi07174/Mi13910
ii
ns
ns
Average No of NBs/brain lobe
150
100
50
0
y*w*;Insc>GAL4
Insc>GAL4; UAS-CSN1b RNAi (RSN2)
Insc>GAL4; UAS-CSN1b RNAi (RSN3)
B
y*w*
y*w*; Mi13910/CSN7T2A>GAL4
y*w*; Mi07174/Mi13910
Dlg Dpn DCP-1
i
ii
iii
C
y*w*; Insc>GAL4
Insc>GAL4;
CSN1b RNAi (RSN2)
Insc>GAL4;
CSN1b RNAi (RSN3)
Mira Dpn DCP-1
i
ii
iii

**Figure S2: *CSN1b* and *CSN7* deficient NBs do not undergo apoptosis.** (A) i, quantification of average NB numbers in larval central brains (72 hours ALH) from y**w** control (n=13), *CSN7* mutants *y*w*; Mi13910/CSN7*$^{T2A>GAL4}$ (n=13) and *y*w*; Mi07174/Mi13910* (n=14), ii quantification of average NB numbers in third instar larval central brains in *y*w*; Insc>GAL4, UAS-mCD8::GFP* control (n=22) and *CSN1b* RNAi mediated knock down (RSN2, n= 24 and RSN-3, n= 23). Error bars represent SEM. n.s. = not significant (Kruskal-Wallis test followed by Dunn's multiple comparison test). (n=number of brains) (B) third instar larval brains from *y*w** control (i, n=6), *CSN7* mutants *y*w*; Mi13910/CSN7*$^{T2A>GAL4}$ (ii, n=6) and *y*w*; Mi07174/Mi13910* (iii, n=7) stained with anti-Dlg (green), anti-Dpn (red), and anti-DCP-1 (cyan), (n=number of brains). Scale bars = 50 µm (C) Mature L3 Larval brains from *y*w*; Insc>GAL4, UAS-mCD8::GFP* control (i, n=10) and *CSN1b* RNAi mediated knock down (RSN2, ii, n= 9 and RSN3, iii, n= 12) stained with anti-Mira (green), anti-Dpn (red), and anti-DCP-1 (cyan), white arrow shows NB positive for DCP-1, (n=number of brains). Scale bars = 50 µm

### Figure S3

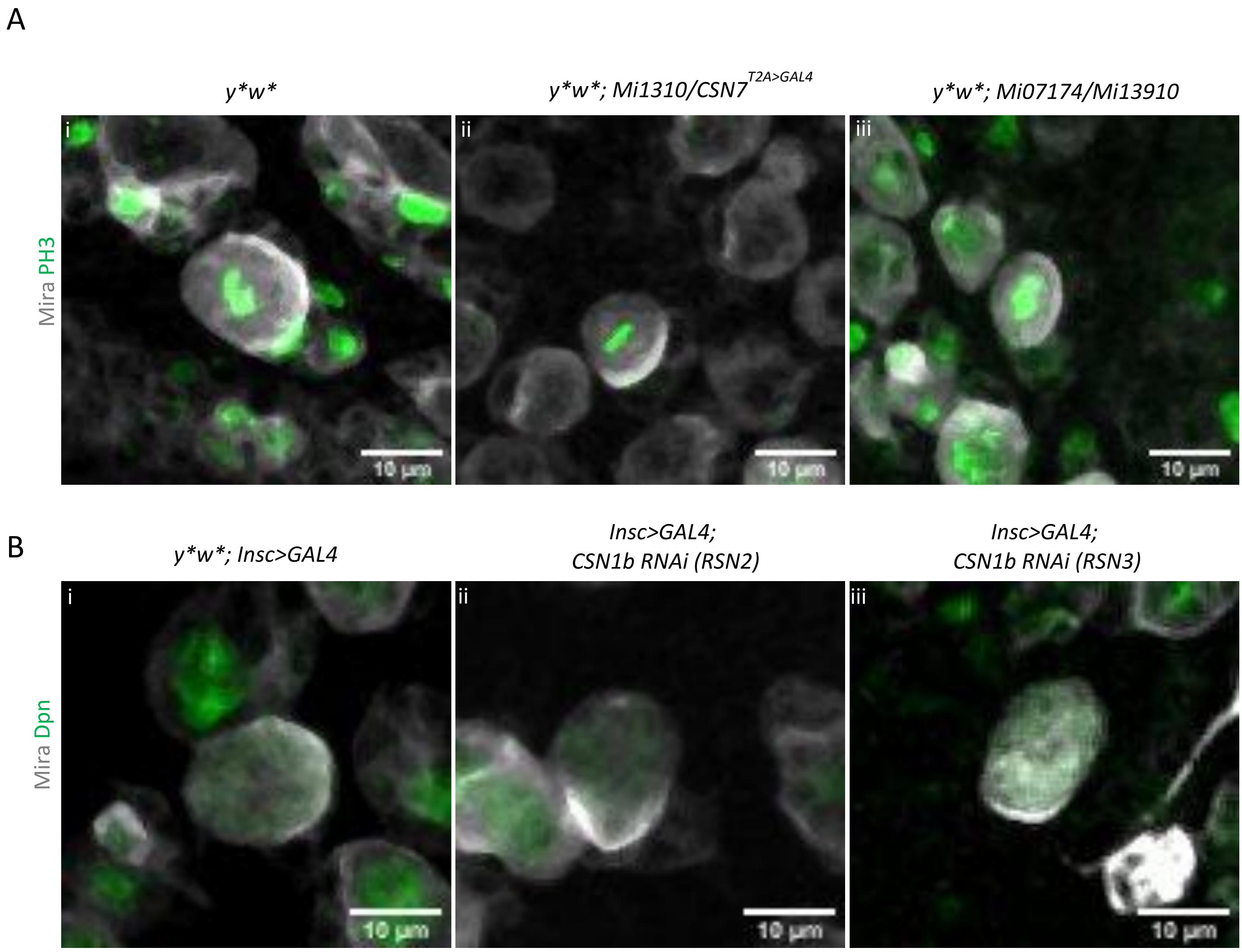


**Figure S3: Loss of *CSN7* and *CSN1b* doesn't affect NB polarity**. (A) High-magnification confocal images of 72 hours ALH larval brains NBs from *y*w** control (i), *CSN7* mutants *y*w*; Mi13910/CSN7*$^{T2A>GAL4}$ (ii) and *y*w*; Mi07174/Mi13910* (iii) undergoing mitosis. Brains were stained for the mitotic marker phosphohistone H3 (PH3, green) and the basal cortical marker Miranda (Mira, gray). Scale bars = 10 µm (B) High-magnification confocal micrographs of mitotic NB from *y*w*; Insc>GAL4, UAS-mCD8::GFP* control (i) and *CSN1b* RNAi mediated knock down (RSN2, ii and RSN3, iii) stained with anti-Mira (gray) and anti-Dpn (green). Scale bars = 10 µm

**Figure S4**

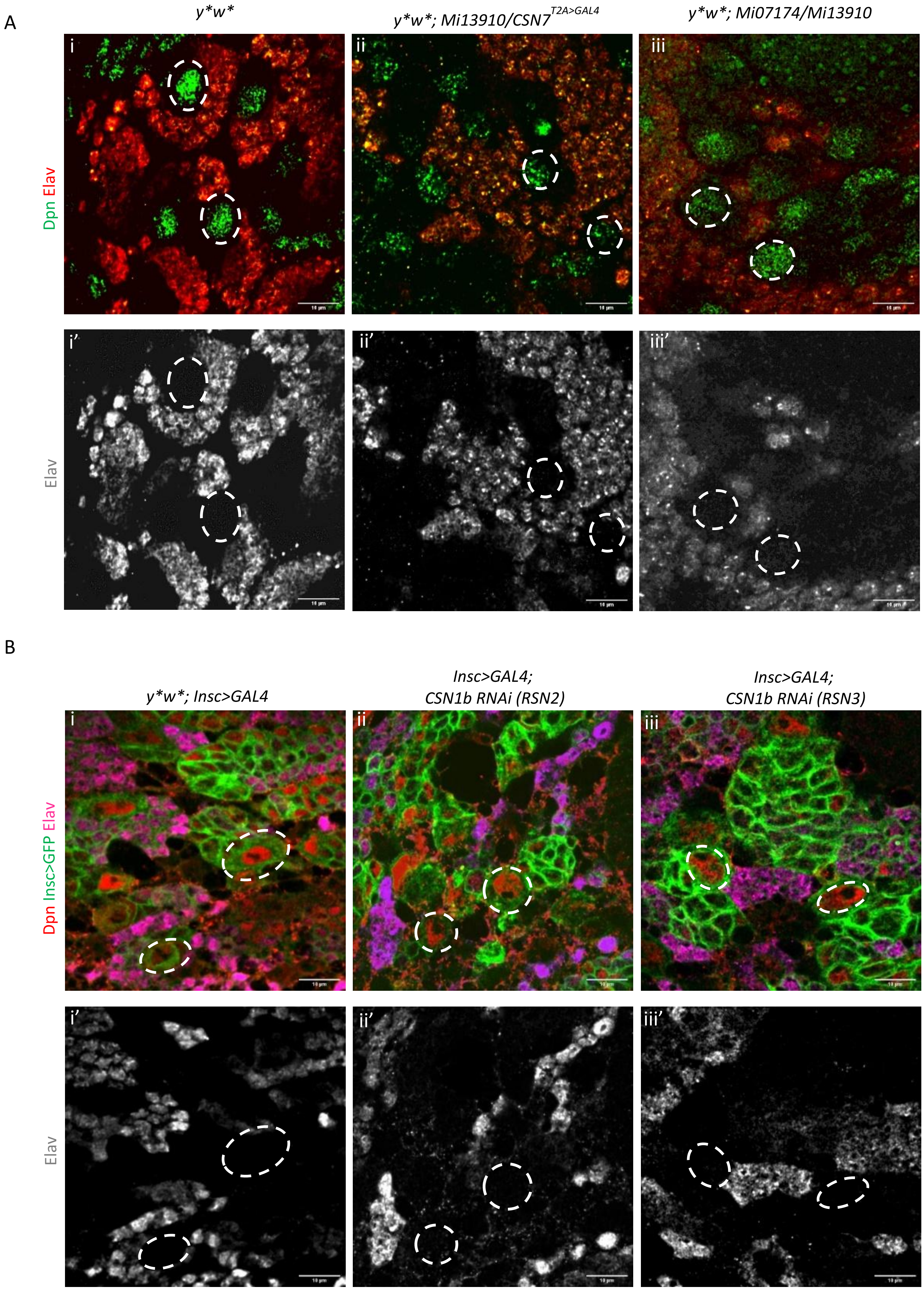
A
y*w*
y*w*; Mi13910/CSN7T2A>GAL4
y*w*; Mi07174/Mi13910
i
ii
iii
i'
ii'
iii'
Dpn Elav
Elav
B
y*w*; Insc>GAL4
Insc>GAL4;
CSN1b RNAi (RSN2)
Insc>GAL4;
CSN1b RNAi (RSN3)
i
ii
iii
i'
ii'
iii'
Dpn Insc>GFP Elav
Elav

**Figure S4: Loss of *CSN7* or *CSN1b* doesn't cause premature differentiation into neurons.** (A) 72 hours ALH larval brains from *y*w** control (n=12), *CSN7* mutants *y*w*; Mi13910/CSN7*$^{T2A>GAL4}$ (n=13) and *y*w*; Mi07174/Mi13910* (n=16) stained with anti-Dpn (red) and anti-Elav (red and gray bottom row). Dpn-positive neuroblasts (gray circles) do not express Elav. Scale bars = 10 µm (B) Third-instar larval brains from *y*w*; Insc>GAL4, UAS-mCD8::GFP* control (n=10) and *CSN1b* RNAi mediated knock down (RSN2, n= 12 and RSN3, n= 9) stained with anti-Dpn (red) and anti-Elav (magenta and gray bottom row) neuroblasts marked by white circles are negative for Elav. Scale bars = 10 µm

**Figure S5**

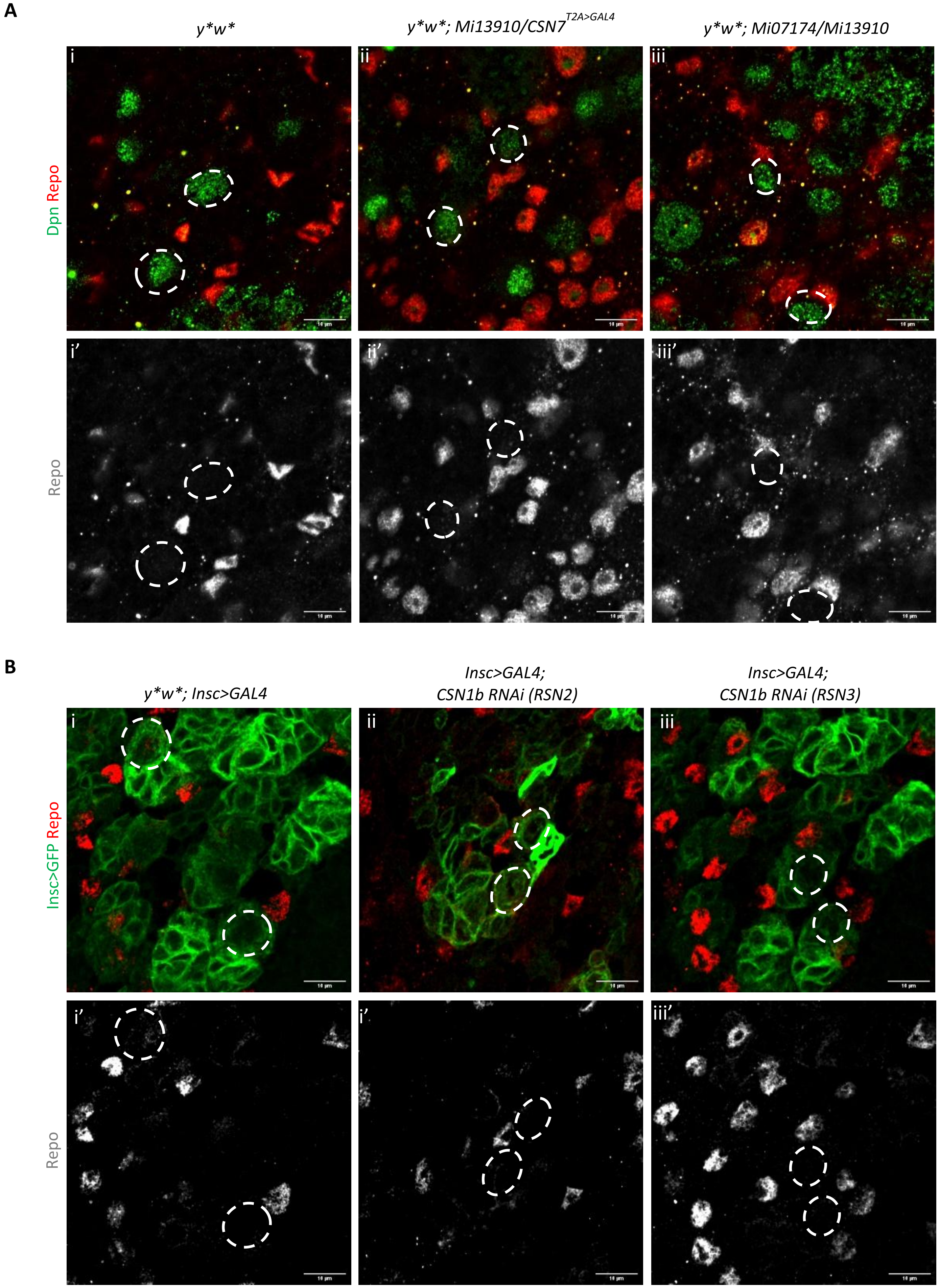
A
y*w*
y*w*; Mi13910/CSN7T2A>GAL4
y*w*; Mi07174/Mi13910
i
ii
iii
Dpn Repo
i'
ii'
iii'
Repo
B
y*w*; Insc>GAL4
Insc>GAL4; CSN1b RNAi (RSN2)
Insc>GAL4; CSN1b RNAi (RSN3)
i
ii
iii
Insc>GFP Repo
i'
i'
iii'
Repo

**Figure S5: Absence of Repo misexpression in NBs suggests no premature differentiation into glia.** (A) 72 hours ALH larval brains from *y*w** control (n=7), *CSN7* mutants *y*w*; Mi13910/CSN7*$^{T2A>GAL4}$ (n=10) and *y*w*; Mi07174/Mi13910* (n=10) stained with anti-Dpn (red) and anti-Repo (red and gray bottom row). Dpn-positive neuroblasts (white circles) do not express Repo. Scale bars = 10 µm (B) 72 hours ALH larval brains from *y*w*; Insc>GAL4, UAS-mCD8::GFP* control (n=8) and *CSN1b* RNAi mediated knock down (RSN2, n=12 and RSN3, n=14) stained with anti-GFP (green) and anti-Repo (Red and gray bottom row) neuroblasts marked by white circles are negative for Repo. Scale bars = 10 µm

### Figure S6

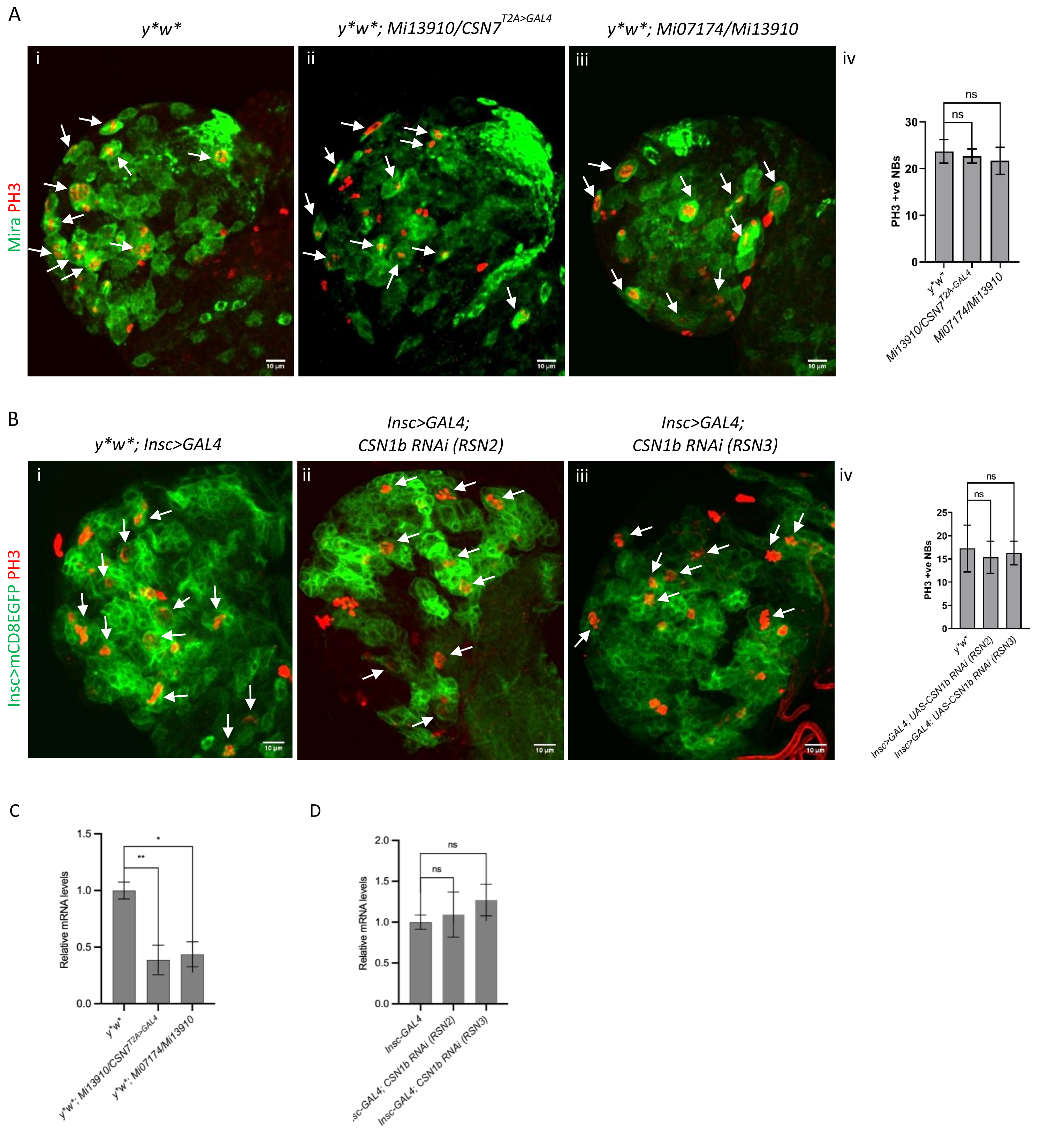

A
y*w*
y*w*; Mi13910/CSN7T2A>GAL4
y*w*; Mi07174/Mi13910
i
ii
iii
iv
Mira PH3
10 μm
PH3 +ve NBs
ns
ns
30
20
10
0
y*w*
Mi13910/CSN7T2A-GAL4
Mi07174/Mi13910
B
y*w*; Insc>GAL4
Insc>GAL4; CSN1b RNAi (RSN2)
Insc>GAL4; CSN1b RNAi (RSN3)
Insc>mCD8EGFP PH3
PH3 +ve NBs
25
20
15
10
5
0
y*w*
Insc>GAL4; UAS-CSN1b RNAi (RSN2)
Insc>GAL4; UAS-CSN1b RNAi (RSN3)
C
Relative mRNA levels
1.5
1.0
0.5
0.0
**
*
y*w*
y*w*; Mi13910/CSN7T2A>GAL4
y*w*; Mi07174/Mi13910
D
Relative mRNA levels
2.0
1.5
1.0
0.5
0.0
ns
ns
Insc-GAL4
Insc-GAL4; CSN1b RNAi (RSN2)
Insc-GAL4; CSN1b RNAi (RSN3)

**Figure S6: *CSN7* and *CSN1b* deficient neuroblasts exit quiescence and enter into mitosis** (A) 24 hours ALH larval brains from *y*w** control (i, n=9), *CSN7* mutants *y*w*; Mi13910/CSN7*$^{T2A>GAL4}$ (ii, n=4) and *y*w*; Mi07174/Mi13910* (iii, n=9) stained with anti-Mira (green) and anti-PH3 (mitotic marker, blue) antibodies. White arrows mark PH3 positive NBs, Controls and *CSN7* mutants show a comparable number of mitotic neuroblasts at 24h ALH. Scale bars = 10 µm. Quantification of the number of PH3-positive neuroblasts per brain lobe is shown in iv. Statistical significance was determined using a Kruskal-Wallis test followed by Dunn's multiple comparison test (ns = not significant, $p > 0.05$) (B) 24 ALH larval brains from *y*w*; Insc>GAL4, UAS-mCD8::GFP* control (n=18) and *CSN1b* RNAi mediated knock down (RSN2, n= 14 and RSN3, n= 14) stained with anti-GFP (green) and anti-PH3 (Red). White arrows mark PH3 positive NBs, controls and *CSN1b* deficient brains show a comparable number of mitotic neuroblasts at 24h ALH. Scale bars = 10 µm. Quantification of the number of PH3-positive neuroblasts per brain lobe is shown in iv. Statistical significance was determined using a Kruskal-Wallis test followed by Dunn's multiple comparison test (ns = not significant, $p > 0.05$) (C) shows quantification of Ago mRNA levels in 72 hours ALH larval brains from *y*w** control, *CSN7* mutants *y*w*; Mi13910/CSN7*$^{T2A>GAL4}$ and *y*w*; Mi07174/Mi13910* (n=3). Statistic test (*$p<0.05$,**$p<0.01$, n.s. = not significant) (D) shows quantification of Ago mRNA levels in third-instar larval brains from *y*w*; Insc>GAL4, UAS-mCD8::GFP* control and *CSN1b* RNAi mediated knock down (RSN2 and RSN3) (n=3). Statistic test (n.s. = not significant)

**Figure S7**

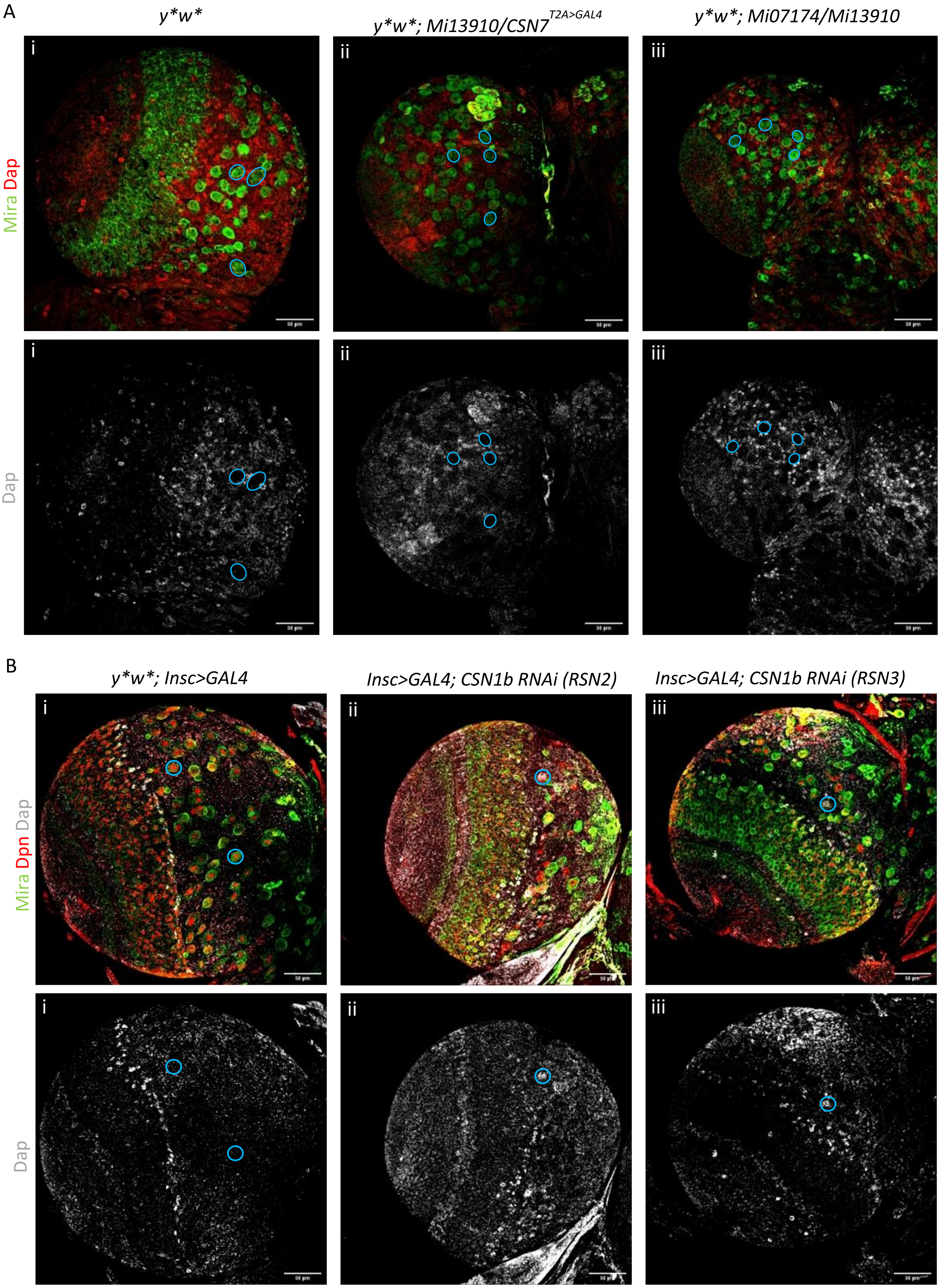

A
y*w*
y*w*; Mi13910/CSN7T2A>GAL4
y*w*; Mi07174/Mi13910
Mira Dap
Dap
i
ii
iii
B
y*w*; Insc>GAL4
Insc>GAL4; CSN1b RNAi (RSN2)
Insc>GAL4; CSN1b RNAi (RSN3)
Mira Dpn Dap
Dap

**Figure S7: *CSN7* and *CSN1b* deficient NBs do not accumulate Dap/p21**. (A) 72 hours ALH larval brains from *y*w** control (i, n=4), *CSN7* mutants *y*w*; Mi13910/*$CSN7^{T2A>GAL4}$ (ii, n=7) and *y*w*; Mi07174/Mi13910* (iii, n=6) stained with anti-Mira (green), anti-DAP (red, gray bottom row) antibodies. Blue circles mark NBs, Mira-positive neuroblasts (blue circles) do not express DAP. Scale bars = 50 µm. (B) Third-instar larval brains from *y*w*; Insc>GAL4, UAS-mCD8::GFP* control (n=5) and *CSN1b* RNAi mediated knock down (RSN2, n= 6 and RSN3, n= 6) stained with anti-GFP (green), anti-Dpn (Red) and anti-DAP (gray, gray bottom row) antibodies. Blue circles indicate NBs with DAP accumulation. Scale bar = 50 µm

**Figure S8**

A B

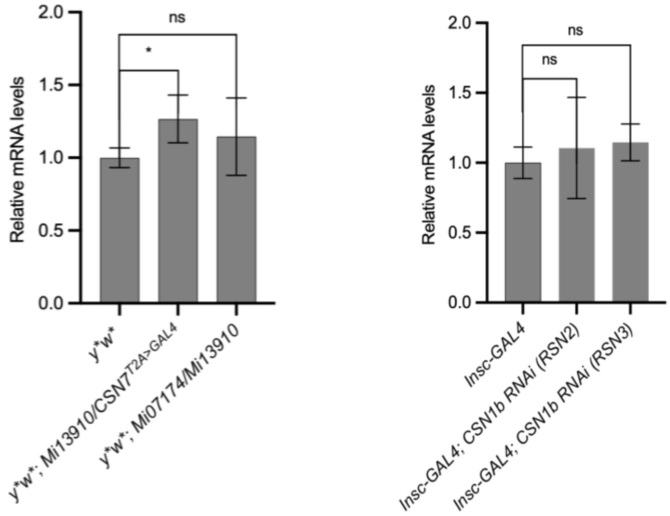


**Figure S8: *CSN7* and *CSN1b* loss doesn't affect CycE expression.** (A) shows quantification of CycE mRNA levels in 72 hours ALH larval brains from *y*w** control, *CSN7* mutants *y*w*; Mi13910/CSN7*$^{T2A>GAL4}$ and *y*w*; Mi07174/Mi13910* (n=3). Statistic test (*p<0.05, n.s. = not significant) (B) shows quantification of CycE mRNA levels in third-instar larval brains from *y*w*; Insc>GAL4, UAS-mCD8::GFP* control and *CSN1b* RNAi mediated knock down (RSN2 and RSN3) (n=3). Statistic test (n.s. = not significant)

**Figure**

A B

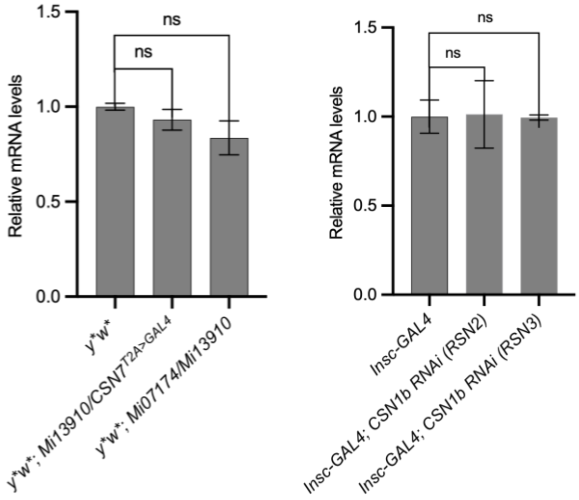


**Figure S9: *CSN7* and *CSN1b* loss doesn't affect Akt mRNA levels.** (A) shows quantification of Akt mRNA levels in 72 hours ALH larval brains from *y*w** control, *CSN7* mutants *y*w*; Mi13910/CSN7*$^{T2A>GAL4}$ and *y*w*; Mi07174/Mi13910* (n=3). Statistic test (n.s. = not significant) (D) shows quantification of Akt mRNA levels in third-instar larval brains from *y*w*; Insc>GAL4, UAS-mCD8::GFP* control and *CSN1b* RNAi mediated knock down (RSN2 and RSN3) (n=3). Statistic test (n.s. = not significant)
